# Interaction-finder: automated literature-based discovery of biological entity associations with quote-level provenance

**DOI:** 10.64898/2026.07.07.736901

**Authors:** Timothy E. Chapman, Timo Laßmann

**Affiliations:** The Kids Research Institute Australia

## Abstract

Identifying interactions between biological entities is a cornerstone of molecular research, but assembling such lists from the literature is slow and tedious. For many research questions, no curated database exists, leaving researchers to survey the relevant literature themselves. We present interaction-finder, a tool that automates this process: given a topic string and user-defined entity types, it discovers relevant literature through llm-guided iterative search, extracts candidate associations from full-text articles, and produces a ranked list where every association is backed by quoted passages verified against the source text. A self-contained interactive html report enables rapid triage of the results. Evaluated across 60 topics in three domains (celltype–cellmarker, disease–gene, and ligand–receptor), interaction-finder recalls 1.2–4.3× as many known associations as single-shot prompting and an off-the-shelf deep-research framework, with all extracted quotes verified against source text.

To assess candidates unrecognised from the gold-standard databases, we scored each candidate using an independent llm judge blind to the tool’s reasoning. Across the three domains, unverified candidates score similarly to gold-standard associations. We find the gold-standard associations are enriched at the top of our ranked candidates, with an overall recall@20 of 0.61.

Interaction-finder is freely available at https://github.com/tecosaur/interaction_finder under an mit licence.

## 1 Introduction

Studying a biological research question often involves considering the interactions between specific entities, such as *which genes are associated with episodic paroxysmal anxiety?* or *what receptors interact with vegfb?* For well-studied relationship categories, such as disease–gene links, researchers can access large curated datasets such as omim [1], DisGenet [2], and ClinVar [3], with smaller datasets available for more specialised topics [4]. When no relevant dataset exists for a research question, or when a researcher needs up-to-date information, manual literature review is required. However, given the rapid growth in the number of manuscripts being published, obtaining a comprehensive overview has become challenging even for domain experts.

Literature mining systematically retrieves information from a larger swathe of the literature than a researcher can hope to read. The idea of extracting relationships from literature was first articulated before automation was feasible in Swanson’s seminal work discovering relationships across papers [5]. This created a framework known as “literature based discovery” (lbd), inspiring computational systems such as bitola [6] that automate the approach using statistical term co-occurrence. Classical literature-based discovery systems excel at generating hypotheses but sacrifice granularity: they produce ranked associations and operate on aggregated co-occurrence statistics rather than specific extracted claims [7].

Natural language processing enables the extraction of relationships between entities described in a paper, such as gene–gene regulatory relationships or enzyme–substrate interactions. A large number of general and specialised tools are now available, which can be broadly organised into families (Table 1). Pre-indexed annotation tools like PubTator 3 [8] make mined information available at scale by processing all PubMed abstracts. Full-text mining consistently recovers more biological associations [9], but operating on abstracts alone is common as they are more readily available. PubTator 3 offers easy and fast entity lookups, but only for the six entity kinds recognised by the system. Other extraction systems such as Bionext [10] offer end-to-end extraction and classification of relationships for arbitrary entity types, but require annotated training data for the specific entities and relationships.

**Table 1:** Comparison of approaches to biological entity-association discovery. An expanded 16-tool comparison is in Supplementary Section S7.

| Tool category | Representative tools | Entity types | Query mode | Discovery | Evidence |
| --- | --- | --- | --- | --- | --- |
| Curated databases | OMIM, DisGeNET, ClinVar | Gene–disease | Lookup | No | Reference |
| Pre-indexed annotation | PubTator 3, BERN2 | Fixed (6–9) | Query | No | Document |
| Classical LBD | ARROWSMITH, BITOLA | MESH terms | Interactive | Yes | Co-occurrence |
| Literature-scale mining | SciLinker | Fixed (4) | Batch | No | Document |
| Supervised RE | BioNext | Fixed schema | Documents | No | Document |
| Document-level LLM RE | SPIRES, SYRACT, LLM-IE | Custom | Documents | No | Sentence |
| Query-driven LLM RE | FuncFetch | Domain-specific | PubMed query | Partial | Document |
| LBD + LLM | SKIM-GPT, LORE | User terms | Interactive | Yes | RAG chunks |
| **On-demand discovery** | **interaction-finder** | **User-defined** | **Topic string** | **Yes** | **Validated quotes** |

Large language models (llms) have substantially expanded what is possible through literature mining. Recent efforts have combined classical literature-based discovery techniques with llms [11,12]. llm-based extraction frameworks like spires [13], Syract [14], and llm-ie [15] support custom entity types, but require researchers to discover relevant literature and assemble the input corpus themselves. FuncFetch [16] extends this pattern by accepting a PubMed query and processing full-text manuscripts, but exclusively targets enzyme–substrate relationships. Other researchers have developed approaches that work across a collection of documents, like lore [17] and SciLinker [18], for a single specific target. At the simpler end of the spectrum, many publications form part of the “world knowledge” embedded in llm training data. There are also a number of llm-driven “deep research” workflows [19,20] that perform iterative retrieval and synthesis.

Despite a large number of computational tools automating the extraction of biological relationships from the literature, researchers interested in a novel or poorly studied biological relationship must still read and assemble a corpus of relevant papers themselves.

Beyond this gap in automation, a major challenge with llm-based approaches is verification and confabulation. Even state-of-the-art models are prone to hallucinating citations and factual claims. This led to the early recognition of the value of requiring llms to quote source material as supporting evidence [21]; however, across studies that use llms to generate text with evidence, few use quotes as evidence [22].

To the best of our knowledge, no existing tool combines automatic keyword-driven search, user-defined entity types, full-text processing, and verifiable quote-level provenance, where every reported association is traceable to specific quoted passages from source articles, each verified against the original text. Interaction-finder fills this gap: given a topic and entity types, it discovers relevant literature, extracts candidate associations from full-text articles with validated quotes, and produces a ranked list for expert review.

## 2 Methods

Interaction-finder takes a natural-language description of a research topic and a list of user-provided entity types. It then runs multiple rounds of literature searches, retrieves full-text articles (where available, falling back on abstracts), then extracts, consolidates, and ranks evidence of associations. The final output is an html report containing a ranked list of candidate associations, each backed by verified quotes from source articles. A high-level overview of the design is shown in Fig. 1; a more detailed diagram and description are available in Supplementary Section S1.

**Figure 1:**
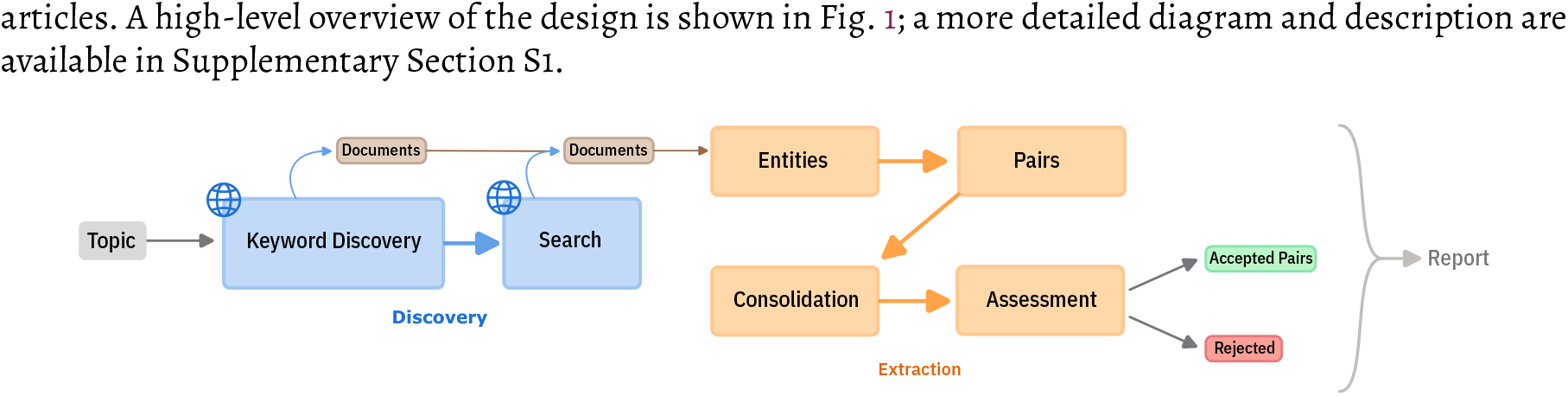
The pipeline. Given a research topic and entity types, the system looks for review articles and extracts keywords from them, uses them to drive multiple rounds of searching, then extracts entities and pairs from the combined results. The complete workflow is shown in Fig. S1.

### 2.1 Iterative llm-guided search

Rather than requiring a pre-assembled corpus, interaction-finder discovers relevant articles directly from a short topic description. Constructing a comprehensive search is non-trivial: the same entity may be known by different names, have subtypes that are individually discussed, or be part of a larger category. Effective search relies on finding the right keywords [23]. For that reason, we split the search into two phases: first, review articles are retrieved and mined for keywords using multiple keyword extraction algorithms (yake, rake, tf-idf, and Keybert), filtered for relevance using an llm. These keywords are combined with the topic to generate a diverse set of queries, which can be run with configurable search backends (PubMed, Searxng, Perplexica, and Openai web search). An llm review step runs up to five rounds of keyword discovery and three rounds of search (both configurable), stopping early when an llm reflector [24] either judges search coverage to be sufficient, or directs further queries toward identified gaps.

### 2.2 Extraction of verified quotes, entities, and pairs from full text

Interaction-finder fetches articles using a web crawler with custom link-following rules, falling back on abstracts when full text is unavailable. Each article is assessed against a paper quality rubric (scoring criteria in Supplementary Section S5), split into chunks [25], and passed to an llm that identifies instances of the user-specified entity kinds with supporting quotes. Since the same entity often appears under different names across papers, entities are consolidated through multiple merging passes [26,27] (see Supplementary Section S4 for the merging approach).

Matching llm-provided quotes to the source text is non-trivial, due to frequent formatting changes and paraphrasing. To address this, we first introduce a consistent reformatting pass that is uniformly applied to both the source text and the llm output. We then use fuzzy text matching with a minimum similarity threshold of 0.75 to locate approximate matches, and use local word-sequence alignment to determine the match candidate. Poor quality matches are inspected to determine the likely failure mode, and used to help the llm retry. See Supplementary Section S2 for the matching approach and error classification. Quotes that fail to match are excluded.

Candidate pairs are identified by grouping entities whose mentions co-occur within a window of two semantic chunks, then asking an llm to assess each co-occurrence. Along with the pair itself and supporting quotes, we ask the llm to provide rubric-based scores of the evidence quality (1–9), and the relevance to the topic (1–5). This information is collected as a ranking signal for the candidates.

All unique pairs across all documents are combined and judged. Pairs with consistent supporting evidence across documents are accepted without llm judgement, using a confidence score computed from the per-document scores. All other pairs, including those with contradictory evidence (e.g., a gene reported as both promoting and suppressing a phenotype), are put to an llm judge to produce a final accept/reject decision, together with a decision confidence and topic relevance score. This provides us with a list of accepted candidates, with supporting judgements and quotations.

To improve the relevance of the candidates presented, we ask an llm to identify specific terms asked about in the topic, and based on that collect a list of extracted entities of the same kind, and ask an llm to assess whether each entity describes the term (is an alias of, is a more specific kind, or a group that it is the major member of), or is merely tangentially related. The verdicts are recorded and used to restrict the associations presented, with the option of viewing the full set. Unless otherwise noted, all statistics reported here are based on the relevance-restricted candidates.

### 2.3 Interactive report for expert triage

Because interaction-finder typically produces tens to hundreds of candidates per topic, the tool generates a self-contained interactive html report that allows a researcher to freely navigate between a sortable list of candidate pairs, the per-document evidence for any pair of interest, and the supporting quotes highlighted in the context of the original article.

### 2.4 Implementation

Interaction-finder is implemented as a Python cli package, installable with `pip` or `uv`. It can be configured at the command line or using a toml configuration file. Language models are used for keyword filtering, entity extraction, pair assessment, and evidence judgement; each stage can be independently assigned to a different model, allowing users to balance cost against capability (see Supplementary Section S11 for the configuration schema). All prompts and output schemas are described in Supplementary Section S14 and can be inspected in the source repository. The keyword generation, search, extraction, and evaluation phases all operate by producing or updating a single json checkpoint file. This enables individual stages to be run or resumed from any checkpoint.

Complete provenance is preserved throughout the pipeline, from search result to final candidate pairs. The full pipeline is detailed in Supplementary Fig. S1.

### 2.5 Evaluation design

The target use case of interaction-finder is to find associations where no curated dataset exists. Since discovery of genuinely novel associations is difficult to evaluate at scale, we instead measure *extraction fidelity*: the pipeline’s ability to recapitulate associations already catalogued in three manually curated and literature-derived gold-standard datasets:

1. Ligand–receptor pairs [28]
2. Cell-types and cellmarkers [29]
3. Disease–gene associations [30]

From these datasets, we selected 60 representative topics: 20 ligands picked at random, then 20 cell types and 20 diseases manually selected for diversity of coverage across physiological systems and reference set sizes (see Supplementary Section S6 for selection criteria).

We compared the associations found by our tool against two basic llm-driven approaches: single-shot prompting (“list all X associated with Y”) and DeerFlow (Deep Exploration and Efficient Research Flow) [31], an off-the-shelf deep research framework. Both DeerFlow and interaction-finder retrieve and synthesise information from the web; single-shot prompting relies on the model’s world knowledge. Single-shot prompting and interaction-finder used gpt-5-mini; DeerFlow used gpt-4.1-mini due to website rate-limiting constraints (see Supplementary Section S6 for a comparison of model effects). Full prompts and configurations are in Supplementary Section S14. Each baseline was run five times per topic to assess robustness. This provides us with a list of gold-standard associations and unverified candidates for each topic, which we evaluate on two axes: the fraction of gold-standard associations the pipeline recovers, and the proportion of its candidates that match a gold-standard entry. While these two axes are analogous to recall and precision, it would be inaccurate to label them as such given the incompleteness of the gold-standard datasets. The gold-standard fraction is a conservative lower bound on the fraction of biologically meaningful associations, rather than an estimate of true precision.

To assess the risk of poor search results and the quality of candidates that fall outside the gold-standard reference, we run two further experiments described below. First, a cross-domain contamination test, which evaluates how robust the pipeline is to irrelevant information. Second, an independent assessment of association plausibility, which tests whether unverified candidates are judged similarly to gold-standard associations. Together, these experiments address the two ways that the pipeline could produce results that are not relevant or biologically meaningful.

### 2.6 Cross-domain contamination test

To evaluate the specificity of the extraction pipeline, we conducted a cross-domain contamination test. Each domain was spiked with documents from a different domain in a circular scheme: disease–gene topics received ligand–receptor articles, celltype–cellmarker received disease–gene, and ligand–receptor received celltype– cellmarker. This ensures that spiked documents contain real biological content (entities, relationships, quotes) but of the wrong kind for the target topic. For each of the 60 topics, we sampled 20 documents uniformly at random from all source-domain topics (with a deterministic per-topic seed for reproducibility), producing contamination fractions of 3–34% of the corpus (14% on average) depending on target corpus size.

We ran per-document extraction on the spiked documents using the same model and settings as the original topic evaluation, then spliced the results into the checkpoint. Spiked entities were not consolidated with the originally extracted entities, to preserve a clean separation between spiked and original results in the analysis. The pipeline was then resumed from the relationship consolidation stage onward, so cross-documentjudgment treated spiked and real assessments identically. We then tracked how many entity pairs and assessments originated from spiked documents at each pipeline stage.

### 2.7 Independent assessment of candidate association plausibility

To estimate what fraction of candidates absent from the reference datasets are genuine associations, we scored each pair’s biological plausibility using an independent llm judge (gpt-5) that had no access to the pipeline’s evidence or reasoning. For each of the 60 topics, pairs were divided into three categories: *gold-standard positives* (accepted pairs matched to the reference dataset), *constructed negatives* (real biological entities from a distant context, unlikely to be genuinely associated; see Supplementary Section S12 for generation details), and *unverified candidates* (accepted pairs absent from the reference, whose status is unknown).

Each pair was presented to gpt-5 with a prompt asking for a direct 1–9 plausibility rating, where 1 indicates no plausible biological connection and 9 indicates a well-established association with mechanistic basis. A small ordinal scale was used, as llms currently produce better judgements on small ordinal scales than binary scores [32] or continuous scales [33]. The order of the two entities within each prompt was randomised to prevent position bias, and each prompt included one positive and one negative calibration example. We selected the direct rating method for the main analysis because it achieved the highest mean auc across domains (0.89) and is the simplest to interpret. Supplementary Section S13 provides prompts, calibration examples, two additional scoring methods that produced results consistent with those presented here, and per-topic score distributions.

To assess the quality of the candidates, we modelled the candidate score distribution as a two-component mixture of the positive and negative score distributions [34]. Each score value defines a category in a multinomial model, and the mixing weight *π* is found by maximum likelihood. We obtained 95% confidence intervals by bootstrapping the labelled positive and negative sets (10,000 resamples).

## 3 Results

We evaluate interaction-finder across three domains (disease–gene, celltype–cellmarker, and ligand–receptor), examining quote verification success rates, the gold-standard associations found, the robustness to irrelevant literature, and whether the candidates absent from the gold standard are judged similarly to the gold-standards in the independent plausibility assessment. We also describe the interactive report used to triage results.

Across the evaluated topics, interaction-finder typically retrieved 50–500 articles per topic, processing approximately ten thousand articles in total (43% full-text, remainder abstracts; see Supplementary Section S3 for per-topic retrieval yield).

### 3.1 Quote verification and llm selection

Every association reported by interaction-finder is traceable to specific quoted passages, each verified against the source text. Over the 60 topics we ran interaction-finder on, the llm extracted 196,000 supporting quotes. Our text matching process located 98% of the quotes within the source text (97–99% per topic), with the remainder flagged as hallucinations and removed from the pipeline. Only quotes matched to a span of the source text were kept and used in the rest of the pipeline.

To investigate the importance of model selection, we ran one topic (“cell-markers for intestinal stem cells”) with gpt-5-nano, gpt-5-mini, and gpt-5, which produced quote match failure rates of 13%, 2%, and 1% respectively. This is consistent with previous findings that hallucination rates increase with decreasing model size [35].

Comparing against our gold-standard dataset, we find that the gold-standard recovery using gpt-5-nano, 44%, is distinctly lower than gpt-5-mini and gpt-5 (78% and 72% respectively). While we see similar performance in terms of quote verification rate and gold-standard recovery using gpt-5-mini and gpt-5, there is an 85× difference in cost ($63 for gpt-5 vs. $0.74 for gpt-5-mini), suggesting that models of a similar size and capability to gpt-5-mini currently make the best cost/capability tradeoff for this task.

Since llms are stochastic machines, they will not produce identical outputs every run. We picked one representative topic from each domain and re-ran it five times. The standard deviation of recall ranged from 5% to 17%, with much greater variation between topics than within topic re-runs (full per-run results are in Supplementary Section S6).

### 3.2 Gold-standard extraction performance

We ran interaction-finder on 20 topics in three domains (celltype–cellmarker, disease–gene, and ligand– receptor), each of which has an associated gold-standard curated dataset. By comparing the candidate list produced by interaction-finder to the gold-standard datasets, we identified how many of the candidates were part of the gold-standard data.

Interaction-finder recovers around half of known associations in the celltype–cellmarker and ligand–receptor domains (65% and 48% respectively; Table 2), exceeding the best baseline in each domain (28% and 41%). The high variance in ligand–receptor gold-standard recovery (± 40%) reflects the small reference sets for individual ligands (1–8 known receptors): four ligands achieved 100% gold-standard recovery while eight with sparse or unfindable literature scored 0%, producing a bimodal distribution. In the disease–gene domain the reference dataset records a greater number of known pairs per term (82 compared to 3–14 in the other domains), and gold-standard recovery is correspondingly lower at 26%, still substantially above the baselines (5–6%). The per-topic gold-standard recovery rates listed in Table 2 are an unweighted average across all topics; the aggregate recovery over all known pairs (63%, 24%, and 45% respectively) is lower for disease–gene because topics with large reference sets and lower recovery contribute more to the aggregate. These figures reflect the default on-topic view; the relevance restriction removes half the reported associations while retaining 90.9% of the recovered gold standard.

**Table 2:** Evaluation results across the three selected domains. “Known” is the total number of reference associations summed across the 20 topics in each domain. Gold-standard recovery and the gold-standard fraction are calculated per topic against the reference datasets. Baseline values are mean ± standard deviation across 5 runs per topic; interaction-finder values are mean ± standard deviation across the 20 topics in each domain.

| Domain | Method | Known | Gold-standard recovery | Gold-standard fraction | Cost/topic |
| --- | --- | --- | --- | --- | --- |
| Celltype–cellmarker | Single-shot | 274 | 28 ± 2% | 35% | <\$0.1 |
| | DeerFlow | " | 22 ± 1% | 26% | <\$0.1 |
| | interaction-finder | " | 65 ± 13% | 17% | \$4.6 |
| Disease–gene | Single-shot | 1634 | 6 ± 0.3% | 72% | <\$0.1 |
| | DeerFlow | " | 5 ± 0.4% | 55% | <\$0.1 |
| | interaction-finder | " | 26 ± 14% | 31% | \$6.1 |
| Ligand–receptor | Single-shot | 55 | 24 ± 7% | 48% | <\$0.1 |
| | DeerFlow | " | 41 ± 1% | 38% | <\$0.1 |
| | interaction-finder | " | 48 ± 40% | 33% | \$1.9 |

The gold-standard fraction–recovery landscape in Fig. 2a-c shows that while deep research consistently finds more candidate pairs than single-shot, gold-standard recovery only improves for ligand–receptor topics. Despite retrieving more documents, deep-research and single-shot achieve similar gold-standard recovery to each other. Interaction-finder exceeds both baselines in gold-standard recovery across all three domains, benefiting from systematic extraction of associations from the retrieved literature. The llm confidence scores produced during pair judgement weakly discriminated the gold-standard associations (see Supplementary Section S8.4 for score distributions), consistent with the poor calibration of llm confidence scores observed in other biomedical nlp tasks [36,37]. To rank the documents, we first examined a document-count based ranking, which achieved a recall@20 of 0.63, but when applied to only recent gold-standard pairs, the recall@20 performance dropped to 0.41 for post-2020 pairs. Because the goal is discovery, the default ranking should favour these recent, less-established associations. Weighting the supporting-document sum by recency makes this explicit trade: it gives up a little overall recall (0.61 versus 0.63) but improves post-2020 recall@20 to 0.57. This recency-weighted sum is the report’s default ordering. This ranking’s enrichment of gold-standard associations is shown in Fig. 2d. See Supplementary Section S9 for more information on the development and evaluation of the ranking.

**Figure 2:**
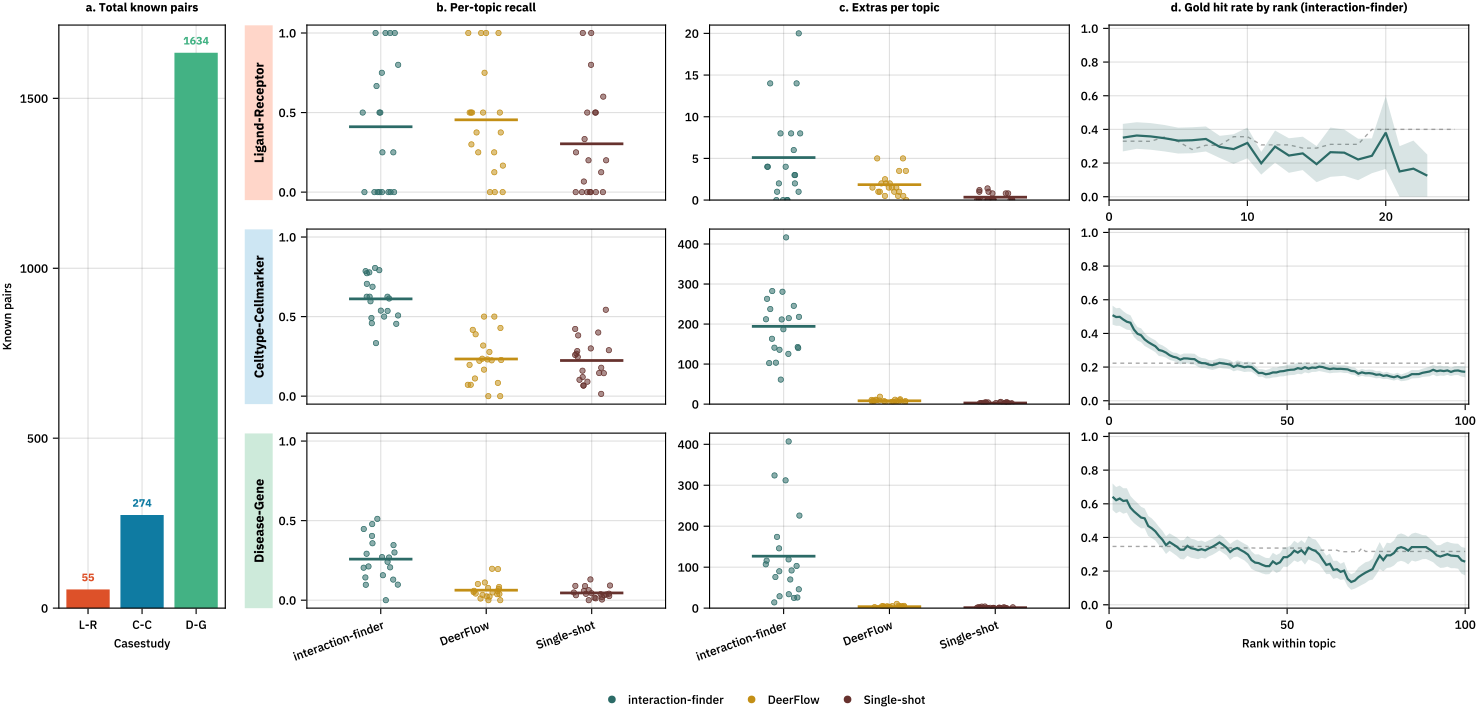
Performance comparison across three domains. **(a)** The number of known pairs in each domain, **(b)** Per-domain recall of gold-standard associations, as a jittered scatter of per-topic values with a horizontal line showing mean recall, **(c)** Per-domain counts of accepted pairs not in the gold-standard (“extras”), shown in the same way; note the differing per-domain y-scales, **(d)** Smoothed gold-standard hit rate at each within-topic rank for interaction-finder, with the dashed line indicating the rank-conditional uniform-ranking baseline; mean ± standard error across topics.

### 3.3 Resilience to irrelevant information

We directly tested the specificity of the pipeline. We expect the biomedical articles retrieved to mention real biological relationships; an indiscriminate pipeline would be liable to extract and present irrelevant relationships. To test this, we injected 20 papers from a different domain into each topic, extracted pairs from them, and combined the results with the original extraction as though the injected papers had been found during search.

Depending on the number of papers originally used, the 20 injected papers constituted 3–34% of the document pool (see Supplementary Section S10 for per-domain and per-topic breakdowns). Across all 1200 injected documents, only 6.5% yielded any entities (compared to 68.5% of real documents), 5.0% produced co-occurring entity pairs, and 3.3% resulted in assessed pairs. So, while the injected documents comprised 10.2% of all documents processed, the 111 pairs that came from them are only 0.15% of the 76,316 assessed pairs from the real documents. This demonstrates a strong ability to filter out irrelevant information.

Of the small number of pairs that did survive, most are genuine associations rather than noise. After cross-document judgement, 9 pairs with weak evidence (score ≤4/9) were rejected, and 99 new pairs accepted from the injected documents. The on-topic gate provides a further line of defence in the report: applied to these 99 pairs it excludes 70 as off-topic. Inspection of the 99 accepted pairs reveals that they are not false positives, but rather a mixture of verified and plausible relationships for the topic in question: 29 match gold-standard verified relations under the same criteria used above. Most of these verified pairs duplicate relations already found from the original documents, which they may have been combined with if not separated out for analysis. Some of the accepted pairs are off topic (filtering for relevance occurs at the search stage), but others directly answer the research question of the topic. As an illustration, a review of proximal tubule endocytosis [38], originally retrieved when searching for receptors that bind to obp2A, was injected into the renal tubular acidosis (rta) disease–gene topic, and yielded three known rta genes (ocrl, ctns, clcn5
) not found by the original search.

### 3.4 Independent plausibility scoring of candidate associations

In the above evaluation, we only considered the retrieval of gold-standard associations. While this is helpful for evaluating against a gold-standard dataset, it leaves us with an incomplete picture of the real relevance of the results. To assess the disposition of the unverified candidates, we constructed negative associations (see Methods), then used three independent llm judges to score the plausibility of the gold-standard, negative, and unverified candidate associations. We used a direct 1–9 score, a 7-point checklist, and a token-probability based score using a different model (gpt-4.1).

We find that the three plausibility scoring methods consistently and reliably separated known positives from constructed negatives. For simplicity, we only discuss the straightforward 1–9 scorer results here, with details on the other two methods placed in Supplementary Section S13. The area under the roc curve for the 1– 9 scorer is 0.84 for disease–gene classification, 0.89 for celltype–cellmarker, and 0.95 for ligand–receptor, confirming that the world knowledge embedded in the llm is sufficient to provide meaningful plausibility scores. This ability to differentiate verified and implausible associations allows us to use the llm judge to assess the unverified candidates.

The score distributions of unverified candidates more closely resembled those of the gold-standard positives than the constructed negative associations (Fig. 3). Modelling each candidate distribution as a mixture of the positive and negative distributions (with bootstrapped 95% confidence intervals), the maximum-likelihood positive-component weights are 79% (ci: 72–83%), 78% (ci: 70–85%), and 98% (ci: 92–100%), for disease– gene, celltype–cellmarker, and ligand–receptor candidates respectively. While the analysis of distributional overlap does not allow us to classify individual candidates, the mixture weights indicate that a large fraction of unverified candidates are judged as similarly plausible to gold-standard positives, suggesting the output merits further inspection.

**Figure 3:**
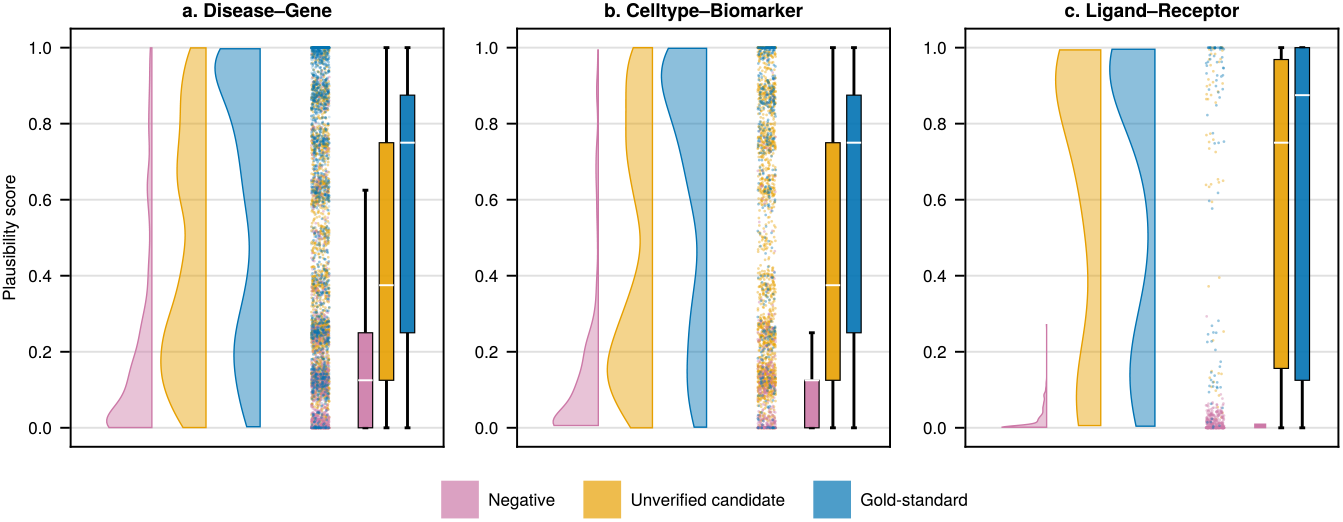
Distribution of plausibility scores across three domains. For each domain, scores are shown for constructed negatives (purple), unverified candidates (orange), and gold-standard positives (blue). Candidates are distributed much more similarly to gold-standard positives than to negatives, indicating that most represent genuine associations absent from the reference dataset.

### 3.5 Interactive report

The interactive html report generated for each topic (Fig. 4) allows a researcher to examine candidate pairs and their underlying evidence. Each topic yields tens to hundreds of candidates, far more than can be evaluated by reading all of the underlying articles; the report makes the focus on gold-standard recovery over strictly matching the reference associations practical through rapid triage. An “On-topic only” switch, enabled by default, restricts the listing to associations whose candidates have been judged as specifically relevant to the topic.

**Figure 4:**
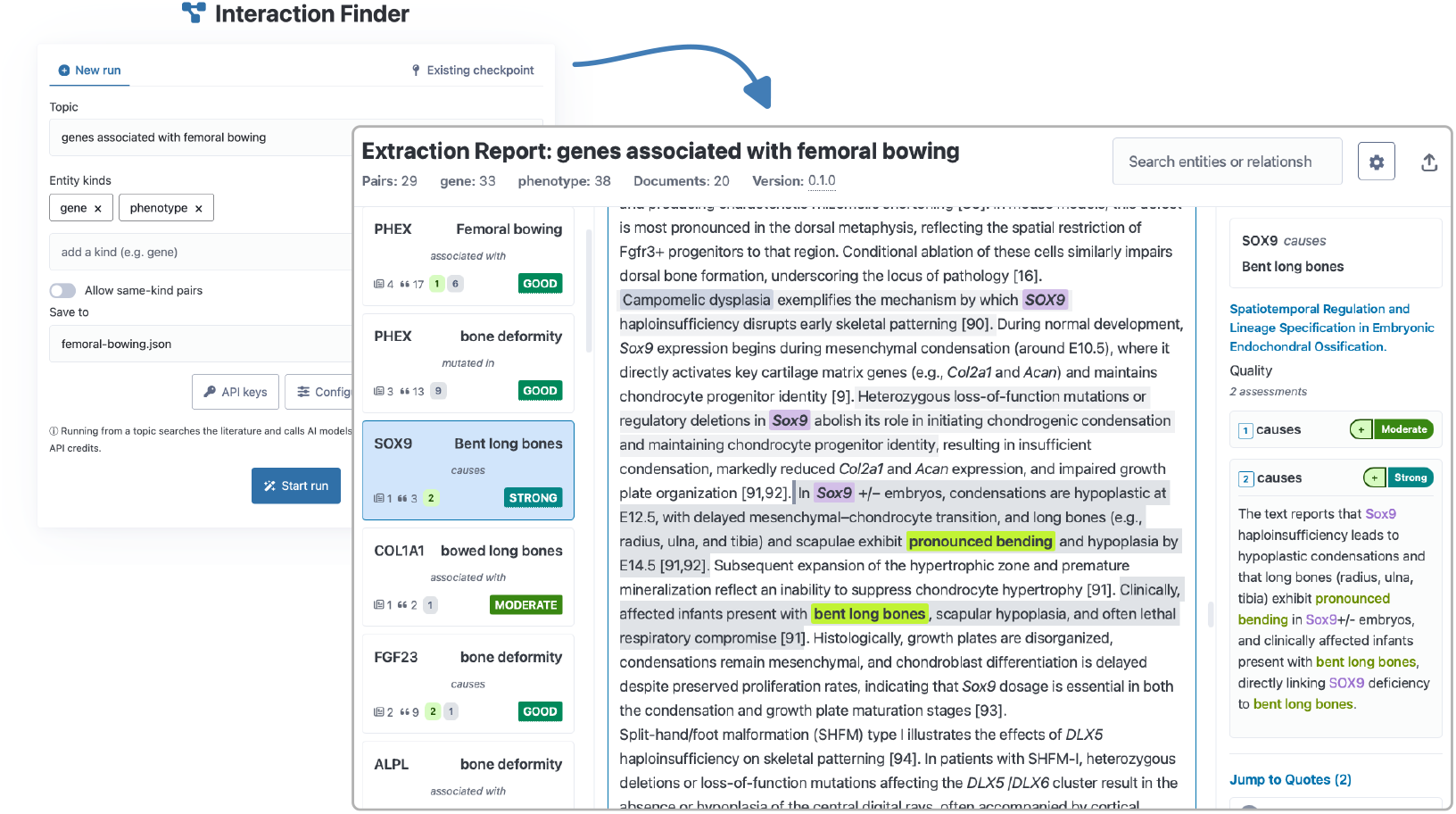
The web interface for generating reports, and a sample report demonstrating the structure of the html report generated by interaction-finder. This self-contained file supports a quick and detailed investigation of the evidence and reasoning behind each result.

Candidate pairs are listed alongside the number of supporting documents and quotes for each, providing an immediate indication ofthe weightofevidence. The candidates are ordered, within the defaulton-topic view, by a recency-weighted sum of supporting documents, which discounts older documents so that recent discoveries are not buried by long-established associations (see Supplementary Section S9 for more information on the ranking). Selecting a candidate displays the source article text with extracted entities and supporting quotes highlighted in context, so that a researcher can verify claims against the original wording. A per-document assessment panel shows the llm’s reasoning chain for each candidate–document pair, making the basis of every judgement transparent. Candidates can be searched, sorted, and filtered; the default ranking serves as a sensible starting point for review, having been found to enrich both gold-standard and higher-plausibility associations. The report is a single html file with no external dependencies, making it straightforward to share with collaborators.

## 4 Discussion

Interaction-finder enables researchers to survey the literature for entity-pair associations; we examined its ability to recapitulate known disease–gene, celltype–cellmarker, and ligand–receptor pairs. However, the approach is entirely generic, and can be extended to domains where no curated dataset exists, such as environmental chemical–epigenetic locus associations, gut microbe–immune cell modulation, or rna modification site–functional consequence mappings. The tool prioritises recall over precision: a missed association is difficult to recover, while a false positive can be quickly identified by reviewing its justification and provenance in the interactive report.

Between 26% and 65% of known associations were recovered across the three domains, with most sources of missed associations external to the extraction pipeline. Full-text access is a major bottleneck: only 43% of retrieved articles were processed as full text, with the remainder falling back to abstracts, which carry less extractable evidence [9]. Incomplete gold-standard recovery is expected: even on curated corpora with fixed entity types, 23% of biomedical relations are missed by the majority of extraction systems [39], and the open-domain setting in which interaction-finder operates is inherently harder. As reference set size grows, gold-standard recovery drops: topics with large numbers of known associations (hundreds in the disease–gene domain) cannot be covered by on-the-fly literature discovery alone. These factors suggest more narrowly focused domains with accessible full-text literature will benefit most from the tool. Beyond coverage, some gold-standard mappings trace indirect ontological relationships (e.g., a disease linked to a gene through a parent phenotype) that the literature does not state directly (see Supplementary Section S8.2 for an illustrative case), and some associations may be documented only in papers not closely related to the query, placing them outside the reach of targeted search. Within the pipeline, entity consolidation is a potential source of both missed and spurious associations: over-merging distinct entities creates false pairs, while failing to merge synonyms splits evidence across duplicate entries.

The gold-standard fractions we reported function as a very conservative analogue of precision, as genuine associations absent from the reference lower this score [40,41]. We estimated this distortion across three domains using a mixture model [34] and found it to be substantial: 78–98% of candidates match the plausibility score distribution of gold-standard associations (higher for domains with smaller reference sets; per-domain estimates and confidence intervals are in the Results). Terminology mismatches between extracted entities and reference entries (e.g., subtypes or naming variants) further depress the score, though the mixture-model corrected estimate should be more robust: name matching determines whether a pair is classified as a confirmed positive or an unverified candidate, but not its plausibility score, so mismatched pairs that are genuinely associated would still be counted as estimated true positives. The plausibility scoring model (gpt-5) belongs to the same model family as the extraction model (gpt-5-mini), which could introduce correlated errors; however, the scoring model discriminates well between known positives and constructed negatives (auc 0.84–0.95), and a scoring method using gpt-4.1 produced consistent estimates (Supplementary Section S13).

Our finding that llm-assigned confidence scores did not reliably discriminate known positives (gold-standard associations) from other candidates is consistent with known calibration challenges in biomedical nlp [36,37]. Multi-document agreement was a stronger triage signal, as pairs cited by more documents with more supporting quotes were more likely to correspond to known associations. Separately, llm stochasticity introduces run-to-run variation in gold-standard recovery, with standard deviations of 5–17% across repeated runs [42,43]. Several pipeline stages constrain this variation, including quote validation, deterministic entity consolidation, and rubric-based scoring with behavioural anchors [44], and the resulting variability was modest relative to the much larger differences between topics’ difficulty.

The contamination experiment showed that 93.5% of injected cross-domain documents were filtered out during entity extraction before reaching pair assessment, and the pairs that survived were genuine associations (for instance, a ligand–receptor paper that also discusses ldlr’s role in hypercholesterolaemia yielded valid disease–gene pairs when injected into a disease–gene topic). Specificity thus operates at the entity-typing stage rather than requiring topic-level filtering, which means the pipeline does not require high search precision, and noisy or over-inclusive retrieval is acceptable as long as entity extraction is sufficiently accurate, as the contamination results suggest it is for the entity types evaluated here.

The tools discussed in the Introduction each address part of the problem of extracting associations from literature; interaction-finder’s contribution is to automate both literature discovery and association extraction with verified provenance in a single workflow [19,45]. Because the contribution is the integrated pipeline, our baselines test the same end-to-end capability: taking a topic and producing an association list. Existing extraction tools such as spires address a different task, requiring pre-assembled corpora and producing sentence-level rather than quote-level evidence.

Where no curated dataset or specialised extraction tool exists, researchers must otherwise assemble evidence from primary literature themselves: the entire 60-topic evaluation processed approximately ten thousand articles for under $300 in api fees, a scale that would be difficult to replicate by manual review. The principal bottlenecks to higher gold-standard recovery (full-text access and search coverage) are external to the pipeline and should diminish as open-access availability continues to expand [46]. Because interaction-finder may be used with commercial llm apis, long-term reproducibility requires that checkpoints be preserved (check-points for the evaluation are deposited with the data; see Data Availability): all post-search pipeline stages can be re-run from a saved checkpoint. The interactive report makes the gold-standard recovery-over-specificity tradeoff practical, letting a researcher trace any candidate association back to the quoted passages and articles that support it, enabling rapid triage without re-reading the underlying literature. Every reported association links back to specific quoted passages in specific papers, providing the verifiable provenance needed to support expert review.

### Key Points

- Interaction-finder automates the path from a research topic to an evidence-backed list of candidate biological associations, each traceable to verified quotes from source articles.
- Across three domains, extracting associations from discovered literature recovers 1.2–4.3× as many known associations as single-shot prompting and deep-research baselines, with all quotes verified against source text and unmatched quotes removed.
- Using an independent llm-based judge, we find gold-standard and plausible associations enriched at the top of the results list.

## Supporting information

Supplementary Sections

## 5 Data availability

Interaction-finder is available at https://github.com/tecosaur/interaction_finder under an mit licence; an archived release is deposited in Zenodo as https://doi.org/10.5281/zenodo.21231829. The evaluation dataset, comprising pipeline checkpoints for all 60 topics, plausibility scores, constructed negatives, and analysis scripts, are available within the same Zenodo record. The gold-standard reference datasets used for evaluation are third-party resources cited in the text.

## 7 Author contributions

Timothy Chapman (Conceptualization, Methodology, Software, Investigation, Validation, Visualization, Writing— Original Draft), Timo Laßmann (Supervision, Conceptualization, Writing—Review & Editing)

