## Supplementary Sections for "Interaction-finder: automated literature-based discovery of biological entity associations with quote-level provenance"

July 7, 2026

**S15 References** **79**

S22 Roster of LLM agents and shared output schemas in the interaction-finder pipeline. 52

### S1 Pipeline architecture

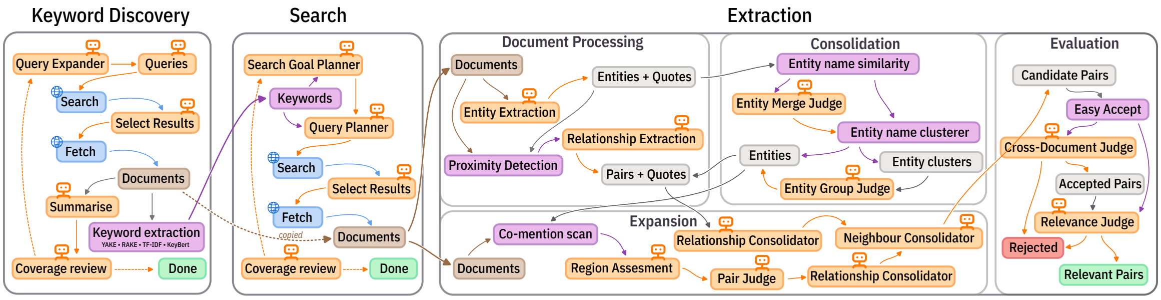

**Figure S1:** Full workflow of the interaction-finder pipeline. The three phases (keyword discovery, search, and extraction) proceed left to right. The extraction phase is further divided into document processing, entity consolidation, pair assessment, and cross-document judgment. Orange rounded boxes denote LLM agent calls; purple rounded boxes denote non-LLM processing steps; rectangular boxes denote data artefacts.

Interaction-finder's pipeline (Fig. S1) comprises three phases (keyword discovery, search, and extraction), each modifying a shared state object and producing a checkpoint. The keyword discovery and search phases both use LLM-driven coverage reviews to decide when to terminate, forming iterative loops rather than fixed-pass processes. The extraction phase first processes each document independently, extracting entities and candidate relationships with supporting quotes. Entities are then consolidated across documents: name-similarity matching proposes merges, an LLM judge confirms or rejects them, and a clustering step groups confirmed aliases. Pair assessment follows a within-then-across-document pattern: relationships are first identified and scored per-document based on textual proximity and co-mention, then a cross-document judge synthesises per-document evidence into a final accept/reject decision for each candidate pair.

#### S1.1 Keyword discovery

The keyword discovery phase (left panel of Fig. S1) identifies "bridging terms" that connect the user's research topic to relevant literature. This echoes the B-terms of Swanson's ABC model (Swanson 1986) but derives them automatically from review articles rather than from manual expert analysis.

The system first searches for review articles on the topic using the configured keyword search backend (SearXNG by default). For each retrieved review, the full text is fetched and converted to Markdown via crawl4ai, then truncated to a configurable character limit (default 50,000 characters). Four keyword extraction algorithms are applied independently to each review, combining three statistical methods with one semantic method:

- **YAKE** (Yet Another Keyword Extractor): statistical keyword extraction based on text features including term frequency, position, and co-occurrence.
- **RAKE** (Rapid Automatic Keyword Extraction): extracts multi-word keywords by identifying phrases delimited by stop words and scoring by word degree and frequency.

- **TF-IDF**: standard term frequency-inverse document frequency scoring using scikit-learn's vectoriser.
- **KeyBERT**: BERT-embedding–based extraction that ranks candidate phrases by their semantic similarity to the document embedding, providing a semantic complement to the statistical methods above.

An LLM agent classifies each candidate keyword as topic-relevant or generic, retaining domain-specific bridging terms. A reflection mechanism then evaluates whether the accumulated keyword set provides sufficient topic coverage, deciding per-review whether to search for additional reviews or conclude.

#### S1.2 Search

The search phase (centre panel of Fig. S1) generates diverse search queries by combining bridging terms with the original topic. Queries are executed across configurable backends:

- **PubMed**: the default backend for biomedical literature, querying NCBI's E-Utilities API.
- **SearXNG**: a meta-search engine aggregating results from multiple web search providers.
- **OpenAI Web Search**: the OpenAI web search API for broader coverage.
- **Perplexica**: a local AI-powered search instance.

Optional semantic reranking using sentence-transformers (model2vec) prioritises results by semantic similarity to the research topic. After each search round, an LLM reflector identifies underexplored aspects of the topic and either directs further queries toward those gaps or terminates the search (Gridach et al. 2025; Huang et al. 2025). Without such validation, LLM-generated search queries can drift from the intended topic (Nazi et al. 2025).

#### S1.3 Entity extraction

The extraction phase (right panel of Fig. S1) processes each discovered article through a multi-step pipeline:

1. **Full-text retrieval.** Articles are fetched via crawl4ai, which handles HTML rendering, JavaScript execution, and PDF extraction. Content is converted to standardised Markdown, enabling consistent downstream processing regardless of source format.
2. **Paper quality assessment.** An LLM agent evaluates each paper using a seven-dimension rubric (Section S5), producing a score from 0–28. Papers scoring below a configurable threshold can be excluded from extraction.
3. **Entity extraction.** Following the structured extraction approach of Dagdelen et al. (2024), an

LLM agent extracts entities of the user-specified types from the full text, along with supporting quotes. Each entity includes its canonical name, any aliases, and the specific text passages where it appears.

4. **Entity consolidation.** Entities are merged across documents through a multi-pass process (Section S4).
5. **Pair identification.** Entity pairs are identified through proximity-based co-mention analysis. Documents are divided into semantic chunks, and entities whose quotes fall within a sliding window of 2 chunks are grouped into "proximal sets." An LLM agent then examines the text region spanning each proximal set and assesses candidate relationships between the co-occurring entities.
6. **Co-mention sweep.** A second pass scans all documents for co-occurrences of entity pairs that were assessed in any document during step 5 but may have been missed in others, for example due to chunking boundaries splitting a co-occurrence, or because entities were not grouped into the same proximal set. For each novel co-mention (i.e. a co-occurrence in a document where that pair has no existing assessment), the surrounding text region is extracted with padding and the pair is assessed by an LLM agent. Overlapping or adjacent regions are merged and assessed in a single call for efficiency.
7. **Evidence assessment.** For each pair in each document, an LLM agent assesses evidence quality using a structured four-dimension rubric (Section S1.5), selects the best relationship type, and identifies supporting quotes.
8. **Relationship consolidation.** Relationship labels are consolidated across documents. Synonymous labels identified by an LLM agent are merged under a canonical label, and each consolidated label is classified by polarity (positive, negative, neutral, or irrelevant). The polarity classification enables detection of "contentious pairs" where contradictory evidence exists, for instance a gene reported as both promoting and suppressing a disease phenotype in different studies.
9. **Cross-document judgment.** A final LLM agent synthesises all per-document evidence for each pair into an accept/reject decision, producing the PairJudgment output that populates the interactive report.
10. **On-topic restriction.** A final agent marks each accepted pair as on- or off-topic (Section S1.4), so that the report can default to the candidates whose subject-side entity genuinely denotes the topic subject. The verdict is recorded on the PairJudgment; nothing is removed.

#### S1.4 On-topic restriction

Each topic has a subject entity, such as a disease, cell type, or ligand, and the pipeline returns candidate pairs naming that subject and a partner. Many of these candidates are off topic: the subject-side entity is not the topic subject but something related to it, such as a downstream consequence, a sibling condition, a derived cell type, or a co-mentioned molecule. The default view should show only the pairs whose subject is genuinely the topic subject, hiding the rest rather than discarding them.

The judgement must not depend on which genes, markers, or partners an entity has: that is the corpus's job, and judging it would import the plausibility call the pipeline keeps separate. The restriction therefore looks only at the taxonomic relation between a candidate and the subject, both of the same kind. One GPT-5-mini call per candidate subject-side entity asks whether the candidate *denotes* the subject (the same entity, a more specific kind of it, a broader category it is a significant kind of, or a grouped label naming it), as opposed to a distinct entity merely related to it. The default verdict is reject.

Two design decisions were load-bearing. First, the subject is passed to the agent *directly* rather than inferred from the topic string. Inference broke ligand–receptor: for "receptors that bind to ADAM2" the model treated receptors as the answer set and rejected ADAM2, the subject itself; supplying `Subject: ADAM2` removes the confusion. The anchor comes from the existing topic-to-anchor extraction, and in a live run it is the search seed. Second, the question asks whether a candidate *denotes* the subject rather than whether it *is* the subject. Asking "is this the subject?" rejects anything non-identical, including legitimate subtypes: it rejected "Parkinson disease" as distinct from parkinsonism rather than a form of it. The *denotes* framing, which admits a taxonomic form of the subject, keeps these subtypes without loosening well-defined subjects.

The verdict is marked on every judgment (a `subject_trust` field, pass or fail), never stripped. The interactive report shows on-topic passes by default and expands to the remainder on a toggle; the same filter flows through to the CLI (`report --filter on_topic:yes|no|unjudged|any`) and the pair export. The restriction's calibration and its effect on the evaluation are reported in Section S6.4.

#### S1.5 Evidence quality scoring

Each per-document evidence assessment decomposes quality into four observable dimensions, listed in Table S1.

**Table S1:** Evidence quality dimensions with levels.

| Dimension | Levels | Assesses |
|---|---|---|
| Directness | explicit, implied, tangential | How directly the source states the relationship |
| Source type | primary, review, other | Type of publication |
| Specificity | mechanistic, associative, vague | Level of detail about the relationship |
| Language | definitive, hedged, speculative | Certainty of the language used |

These dimensions are synthesised into a probability-anchored overall score on a 1–9 scale, where each level corresponds to an approximate probability that the relationship is real: 1 (<5%) through 9 (>95%). Labels range from "None" (1) through "Moderate" (6) to "Robust" (9).

Assessments are grouped by relationship polarity in a PairSpread structure: positive (promoting/activating), negative (inhibiting/protective), neutral (ambiguous directionality), and irrelevant (orthogonal to the research question). This structure enables identification of "contentious pairs" where contradictory evidence exists across documents, for example a gene reported as both promoting and suppressing a disease phenotype.

As reported in the main text, these scores do not meaningfully discriminate true from false positives in our evaluation. Decomposing evidence quality into observable factors should in principle improve calibration. In practice, document count remains a better ranking signal. This is consistent with the poor calibration of LLM self-assessment observed across biomedical NLP tasks (de Oliveira et al. 2025; Li et al. 2023).

### S2 Quote validation system

Quote validation responds to a quantified failure mode: LLMs achieve F1 as low as 0.625 on detecting hard medical hallucinations, with hallucinated content semantically close to ground truth (Pandit et al. 2025). The DAHL benchmark (Seo et al. 2024) further shows that larger models hallucinate less, but with diminishing gains beyond 7–8B parameters. In our evaluation, a single topic re-run with GPT-5-nano showed a reduced quote fidelity of 86.9% compared to 98%+ across the main GPT-5-mini and GPT-5 runs, consistent with this trend.

#### S2.1 Fuzzy matching algorithm

Every LLM-extracted quote is validated against its source document using a position-tracked fuzzy matching system (`NormalizedTextMapper`). The algorithm:

1. **Normalises** both the quote and document text (Unicode normalisation, case folding, whitespace standardisation) while maintaining a position mapping between normalised and original coordinate spaces.
2. **Identifies matching blocks** between the normalised quote and document using sequence alignment, tracking contiguous regions where words align.
3. **Computes similarity** based on the proportion of quote words that align with document text, weighted by contiguity (preferring single contiguous matches over fragmented ones).
4. **Classifies errors** when the match falls below the similarity threshold (default 0.75).

The 0.75 threshold was set by inspecting accept/reject decisions across a manual sweep of candidate values and choosing a conservative cutoff: genuine paraphrases still appear below 0.75, but their alignment to the source becomes increasingly tenuous and harder to verify by eye. The threshold trades a small number of legitimate matches for a sharp reduction in degenerate ones (substring coincidences or unrelated text sharing a few common words), which we judged the safer tradeoff for evidence whose purpose is to be checked by a reader.

#### S2.2 Error classification

When a quote does not exactly match the source document, the system classifies the discrepancy into one of the seven error types defined in Table S2.

Each validated quote receives a correction suggestion containing the best-matching document text, a confidence score (0.0–1.0), and metrics for alignment quality, contiguity, and boundary precision.

**Table S2:** Quote validation error types with descriptions.

| Error type | Description |
|---|---|
| Split quote | Non-contiguous matching blocks (quote spans gap in doc) |
| Word substitution | High overall similarity with specific word mismatches |
| Insertion | Extra words present in quote not in document |
| Deletion | Words missing from quote that are in document |
| Paraphrase | Low similarity but some structural preservation |
| Reordering | Same words present but in different order |
| Not found | No significant alignment with any document passage |

#### S2.3 Validation results

Across the full evaluation (60 topics), 195,684 quotes were validated against source documents with per-domain pass rates of 97.8% (celltype–cellmarker: 86,660/88,612), 98.3% (disease–gene: 89,391/90,894), and 98.7% (ligand–receptor: 19,633/19,897). The lowest individual pass rate (86.9%) was from the intestinal-stem-cell GPT-5-nano run, consistent with the model size effect reported in the hallucination literature. All other runs exceeded 95%.

#### S3 Full-text retrieval yield

Interaction-finder attempts to fetch the full text of every discovered article, falling back to the abstract when full text is unavailable. Because the crawler relies on open-access sources and publisher-permitted HTML, retrieval yield varies by topic and domain. To quantify this, we classified every processed document as full-text or abstract-only using a simple heuristic: documents of at least 5,000 characters with three or more content section headers (markdown headings excluding site metadata such as "Abstract", "Conflict of Interest", and PubMed navigation elements), or exceeding 20,000 characters, were classified as full-text. The length-only threshold captures articles that lose section headings during HTML-to-markdown conversion while retaining their body text. All remaining documents were classified as abstract-only.

Across the 60 evaluated topics, 43% of processed documents (4,500 of 10,550) were classified as full-text (Table S3). Full-text rates varied from 16% (SFTPD, a ligand with sparse literature) to 63% (mature luminal cell), with per-domain averages of 48% (celltype–cellmarker), 42% (disease–gene), and 38% (ligand–receptor). This variation likely reflects differences in open-access availability across research areas. The retrieval yield represents a lower bound on the proportion of full-text articles in the discovered literature, since some articles behind paywalls may have relevant full text that the crawler could not access. Institutional access or integration with licensed full-text sources (e.g. PMC) could improve coverage.

**Table S3:** Full-text retrieval yield per topic. Overall, 43% of processed documents were classified as full-text (range 16–63% across topics), with variation reflecting differences in open-access availability across research areas. Documents of ≥5,000 characters with ≥3 content section headers, or exceeding 20,000 characters, were classified as full-text; all others as abstract-only.

| Domain | Topic | Total | Full | Abstract | Full% |
|---|---|---|---|---|---|
| Celltype-cellmarker | Activated B cell | 152 | 66 | 86 | 43% |
| | Alveolar type II cell | 340 | 204 | 136 | 60% |
| | Capillary cell | 92 | 44 | 48 | 48% |
| | Chondrocyte | 314 | 138 | 176 | 44% |
| | Common lymphoid progenitor cell | 69 | 21 | 48 | 30% |
| | Dopaminergic neuron | 76 | 25 | 50 | 33% |
| | Effector T cell | 114 | 63 | 51 | 55% |
| | Endocardial cell | 104 | 60 | 44 | 58% |
| | Erythroblast | 119 | 62 | 57 | 52% |
| | Hepatoblast | 120 | 46 | 74 | 38% |
| | Intestinal stem cell | 345 | 177 | 168 | 51% |
| | Kupffer cell | 84 | 44 | 39 | 53% |
| | Langerhans cell | 195 | 70 | 125 | 36% |
| | Limbal stem cell | 134 | 68 | 66 | 51% |
| | Mature luminal cell | 79 | 50 | 29 | 63% |
| | Mature neutrophil | 62 | 27 | 35 | 44% |
| | Megakaryocyte progenitor cell | 133 | 51 | 82 | 38% |
| | Neural crest cell | 332 | 155 | 177 | 47% |
| | Plasmablast | 125 | 61 | 64 | 49% |

Continued on next page

Continued from previous page

| Domain | Topic | Total | Full | Abstract | Full% |
|---|---|---|---|---|---|
| | Proximal tubular cell | 116 | 47 | 69 | 41% |
| Disease-gene | Autoimmune thrombocytopenia | 109 | 36 | 73 | 33% |
| | Dilated cardiomyopathy | 367 | 186 | 181 | 51% |
| | Emphysema | 512 | 232 | 279 | 45% |
| | Episodic paroxysmal anxiety | 147 | 50 | 97 | 34% |
| | Femoral bowing | 75 | 36 | 39 | 48% |
| | Hypercholesterolemia | 242 | 108 | 134 | 45% |
| | Hypertriglyceridemia | 154 | 46 | 108 | 30% |
| | Migraine | 274 | 111 | 163 | 41% |
| | Nephrotic syndrome | 381 | 141 | 240 | 37% |
| | Parkinsonism | 330 | 95 | 235 | 29% |
| | Pituitary hypothyroidism | 270 | 87 | 183 | 32% |
| | Progressive hearing impairment | 328 | 123 | 205 | 38% |
| | Psoriasiform dermatitis | 154 | 78 | 75 | 51% |
| | Pulmonary arterial hypertension | 577 | 300 | 277 | 52% |
| | Pulmonary fibrosis | 208 | 99 | 109 | 48% |
| | Renal tubular acidosis | 123 | 28 | 95 | 23% |
| | Rhabdomyolysis | 289 | 114 | 175 | 39% |
| | Rheumatoid arthritis | 398 | 212 | 186 | 53% |
| | Truncal ataxia | 141 | 56 | 85 | 40% |
| | Zonular cataract | 165 | 42 | 123 | 25% |
| Ligand-receptor | ADAM2 | 39 | 7 | 32 | 18% |
| | AREGB | 114 | 62 | 52 | 54% |
| | ASIP | 166 | 43 | 123 | 26% |
| | CGA | 153 | 52 | 101 | 34% |
| | CSF3 | 87 | 38 | 49 | 44% |
| | CXCL6 | 85 | 23 | 62 | 27% |
| | JAG2 | 77 | 46 | 31 | 60% |
| | MFAP2 | 67 | 28 | 39 | 42% |
| | OBP2A | 111 | 39 | 72 | 35% |
| | PDAP1 | 104 | 47 | 57 | 45% |
| | PROS1 | 105 | 50 | 55 | 48% |
| | PSEN1 | 189 | 53 | 136 | 28% |
| | PTHLH | 256 | 103 | 153 | 40% |
| | RAET1E | 40 | 15 | 25 | 38% |
| | SEMA6D | 56 | 17 | 39 | 30% |
| | SERPINA7 | 85 | 24 | 61 | 28% |
| | SFTPD | 45 | 7 | 38 | 16% |
| | SLPI | 74 | 36 | 38 | 49% |
| | TSHB | 180 | 70 | 110 | 39% |
| | VEGFB | 168 | 81 | 86 | 49% |

#### S4 Entity consolidation

A core challenge in extracting entities from multiple documents is the many-names-one-entity problem: the same biological entity may appear as "BMPR2", "bone morphogenetic protein receptor type 2", "BMPRII", or "ALK3" across different papers. Pairwise comparison of all entities is computationally infeasible (quadratic in entity count), so interaction-finder uses a multi-pass hierarchical approach (Dobbins 2024; Fu et al. 2025). The three passes target progressively harder cases at progressively higher cost: deterministic matching catches obvious duplicates without LLM calls, LLM clustering resolves ambiguous name overlaps, and neighbour-based consolidation finds duplicates that only reveal themselves through shared relationship context.

##### S4.1 Pass 1 — automatic fuzzy matching

The first pass identifies obvious duplicates without LLM calls:

- **Capitalisation variants**: entities differing only in case (e.g., "BRCA1" vs "Brca1") are auto-merged, with the mixed-case or conventional form preferred.
- **Parenthetical expansion**: entities where one form is the parenthetical expansion of another (e.g., "SOX9 (SRY-box 9)" and "SOX9") are auto-merged.
- **Token overlap**: entities sharing a high proportion of tokens above a configurable threshold are flagged. Those meeting strict criteria (very high similarity, one clearly a substring of the other) are auto-merged; others are queued for LLM review.

##### S4.2 Pass 2 — LLM-reviewed clustering

Entities that cannot be resolved automatically are grouped into clusters based on token similarity and presented to an LLM agent for decision. The agent receives a cluster of 2+ entity names and decides which should be merged, which should remain separate, and what the canonical name should be.

Clustering uses Reliable Agglomerative Clustering (Chehreghani 2021) with average-linkage similarity and a mutual nearest-neighbour constraint: at each step, only clusters that are each other's best available partner are merged, filtering out fragile merges that would depend on slight ordering differences. Token similarity is weighted by an IDF-like specificity score that accounts for mention frequency and confidence, so rare domain-specific terms (e.g. gene symbols) contribute more than common words. The resulting merge tree can be surgically split by the LLM reviewer when a cluster groups entities that should remain separate. Groups previously resolved are cached to avoid re-review. The process iterates: if merging introduces new canonical names, groups containing those names are re-reviewed until convergence or a maximum iteration count.

This design aligns with the findings of Fu et al. (2025), who report hierarchical cluster merging achieving up to 150% accuracy improvement and 5× fewer API calls than pairwise matching for entity resolution.

##### S4.3 Pass 3 — neighbour-based consolidation

The final pass addresses entities that appear distinct in isolation but are revealed as duplicates by their relationship context. When an anchor entity (e.g., "chondrocyte") connects to multiple similar-sounding neighbours (e.g., "collagen type II" and "COL2A1"), those neighbours may represent the same concept. The algorithm:

1. Builds neighbour sets from pair assessments across all documents.
2. Clusters each anchor's neighbours by token similarity.
3. Presents multi-member clusters to an LLM agent with the anchor context.
4. Applies merge rules to update pair references globally.

This context-dependent approach addresses the homonym disambiguation challenge highlighted by BELHD (Garda and Leser 2024): the same entity name can refer to different concepts in different contexts. By presenting the anchor relationship context, the LLM can distinguish genuine duplicates from coincidental name similarity.

#### S5 Paper quality rubric

Papers retrieved by the search stage are assessed for scientific quality before entity extraction, with the score attached to every claim subsequently extracted from the paper. The score travels with the evidence into the interactive report, letting a reader see at a glance whether a candidate association rests on rigorous or marginal sources, but does not by default gate extraction; a configurable threshold can be set to exclude papers below a chosen quality tier if a researcher wants stricter filtering.

The rubric comprises seven dimensions (Table S4), each scored 0–4, yielding an overall score of 0–28. The dimensions target distinct failure modes in retrieved literature: methodological clarity and statistical rigour assess scientific soundness, data provenance and reproducibility signals assess verifiability, internal consistency and plausibility of claims detect potential errors or overstatement, and integrity indicators flag common red flags (e.g., paper mills, predatory journals). The intent is to exclude low-quality sources without penalising legitimate research that lacks polish.

**Table S4:** Paper quality assessment dimensions with scoring anchors.

| Dimension | 0 (Absent) | 2 (Partial) | 4 (Exemplary) |
|---|---|---|---|
| Methodological clarity | Methods absent | Workflow clear, params missing | Full protocol reproducible |
| Data provenance | Source unknown | Selection partially described | Complete sample flow documented |
| Statistical rigour | No quantitative support | Correct tests, incomplete | Full reporting + validation |
| Internal consistency | Clear contradictions | Generally coherent | Every result traces to method |
| Plausibility of claims | Extraordinary claims | Claims generally supported | Limitations + alternatives named |
| Reproducibility signals | Nothing mentioned | Partial materials | All materials fully specified |
| Integrity indicators | Strong red flags | Neutral | Preregistration, open materials |

Interpretation tiers: 0–7 EXCLUDE (serious deficiencies), 8–14 CAUTION (notable gaps, usable for hypothesis generation), 15–21 ACCEPTABLE (sound methodology), 22–28 HIGH TRUST (rigorous and transparent). The scoring prompt instructs the LLM to treat unqualified perfect metrics (100% accuracy, AUC=1.0) as a yellow flag rather than evidence of high quality, on the grounds that genuinely rigorous papers acknowledge their limitations and unrealistic-looking results often reflect overfitting, leakage, or selective reporting.

### S6 Gold-standard dataset construction

#### S6.1 Ligand–receptor

The reference dataset used for evaluation is the manually curated ligand–receptor interaction dataset from Ramilowski et al. 2015 (Ramilowski et al. 2015), comprising confirmed interactions and 135 manually excluded predictions across 144 human primary cell types. A more recent update, ConnectomeDB2025 (Liu et al. 2026), is now available.

Twenty ligands were selected at random (Table S5). The sample size (n=20) was chosen to support basic statistical characterisation while remaining feasible within the per-topic compute budget. Known receptor counts range from 1 (OBP2A, PDAP1) to 8 (ASIP), totalling 55 reference pairs.

**Table S5:** Selected ligands for the ligand–receptor evaluation domain.

| Ligand | Known receptors |
|---|---|
| ADAM2 | 5 |
| AREGB | 2 |
| ASIP | 8 |
| CGA | 4 |
| CSF3 | 2 |
| CXCL6 | 2 |
| JAG2 | 4 |
| MFAP2 | 1 |
| OBP2A | 1 |
| PDAP1 | 1 |
| PROS1 | 2 |
| PSEN1 | 6 |
| PTHLH | 3 |
| RAET1E | 1 |
| SEMA6D | 4 |
| SERPINA7 | 1 |
| SFTPD | 2 |
| SLPI | 1 |
| TSHB | 1 |
| VEGFB | 4 |

#### S6.2 Celltype–cellmarker

The reference database is CellMarker 2.0 (Hu et al. 2023), containing 83,361 tissue-cell type-marker entries across 2,578 cell types from 656 tissues. The human marker file (`Cell_marker_Human.xlsx`) was obtained from http://www.bio-bigdata.center/CellMarker_download_files/file/Cell_marker_Human.xlsx (SHA-256 `bf52b8cd60df60f1`).

Twenty cell types were selected for diversity (Table S6): 10 blood/immune and 10 other tissues. Known marker counts range from 5 (megakaryocyte progenitor cell) to 35 (neural crest cell), totalling 274 reference pairs.

**Table S6:** Selected cell types for the celltype–cellmarker evaluation domain. The two stem-cell topics are listed at the generic Cell Ontology node CL:0000034 ("stem cell"); marker lookup uses cell-type name rather than CL ID, so this does not affect the gold-standard set.

| Kind | CL family | Name | Known markers |
|---|---|---|---|
| Blood | CL:0002144 | Capillary cell | 14 |
| Blood | CL:0000051 | Common lymphoid progenitor cell | 6 |
| Blood | CL:0000553 | Megakaryocyte progenitor cell | 5 |
| Blood | CL:0000765 | Erythroblast | 9 |
| Blood | CL:0000980 | Plasmablast | 28 |
| Cartilage | CL:0000138 | Chondrocyte | 7 |
| Eye | CL:0000034 | Limbal stem cell | 12 |
| Immune | CL:0000091 | Kupffer cell | 12 |
| Immune | CL:0000096 | Mature neutrophil | 11 |
| Immune | CL:0000236 | Activated B cell | 17 |
| Immune | CL:0000453 | Langerhans cell | 13 |
| Immune | CL:0000911 | Effector T cell | 18 |
| Intestine | CL:0000034 | Intestinal stem cell | 18 |
| Kidney | CL:1000839 | Proximal tubular cell | 20 |
| Liver | CL:0005026 | Hepatoblast | 11 |
| Lung | CL:0002063 | Alveolar type II cell | 9 |
| Misc | CL:0002326 | Mature luminal cell | 6 |
| Muscle | CL:0002350 | Endocardial cell | 12 |
| Neural | CL:0000700 | Dopaminergic neuron | 11 |
| Neural | CL:0011012 | Neural crest cell | 35 |

No clear negative set exists for this domain: very few proteins can be definitively ruled out as cell-markers.

#### S6.3 Disease–gene

The reference is HPO (Human Phenotype Ontology) gene associations from JAX (Jackson Laboratory) (Gargano et al. 2024). The phenotype-to-gene mapping (`phenotype_to_genes.txt`) was taken from the HPO v2025-05-06 release at https://github.com/obophenotype/human-phenotype-ontology/releases/download/v2025-05-06/phenotype_to_genes.txt (SHA-256 `ed68d219529a9d3e`).

Twenty HPO phenotypes were selected for system diversity (Table S7). Gene counts range from 17 (episodic paroxysmal anxiety) to 185 (dilated cardiomyopathy), totalling 1,634 reference pairs.

This domain is notably harder than the other two: reference sets are an order of magnitude larger, and gene-disease associations are often inferred from pathway membership or genetic studies rather than explicitly stated in individual papers.

Table S7: Selected diseases for the disease–gene evaluation domain.

| System | HPO term | HPO ID | Known genes |
|---|---|---|---|
| Cardiovascular | Pulmonary arterial hypertension | HP:0002092 | 164 |
| | Dilated cardiomyopathy | HP:0001644 | 185 |
| Respiratory | Emphysema | HP:0002097 | 63 |
| | Pulmonary fibrosis | HP:0002206 | 78 |
| Endocrine | Hypertriglyceridemia | HP:0002155 | 133 |
| | Pituitary hypothyroidism | HP:0008245 | 38 |
| | Hypercholesterolemia | HP:0003124 | 85 |
| Immune | Autoimmune thrombocytopenia | HP:0001973 | 48 |
| | Rheumatoid arthritis | HP:0001370 | 23 |
| Musculoskeletal | Rhabdomyolysis | HP:0003201 | 45 |
| | Femoral bowing | HP:0002980 | 51 |
| Sensory | Zonular cataract | HP:0010920 | 52 |
| | Progressive hearing impairment | HP:0001730 | 60 |
| Skin | Psoriasiform dermatitis | HP:0003765 | 33 |
| Excretory | Renal tubular acidosis | HP:0001947 | 49 |
| | Nephrotic syndrome | HP:0000100 | 137 |
| Nervous system | Truncal ataxia | HP:0002078 | 77 |
| | Parkinsonism | HP:0001300 | 141 |
| Neurological | Migraine | HP:0002076 | 155 |
| | Episodic paroxysmal anxiety | HP:0000740 | 17 |

#### S6.4 Evaluation methodology

**Terminology.** Following the manuscript, we use *gold-standard recovery* for the fraction of reference associations the pipeline extracts, and *gold-standard fraction* for the proportion of accepted pairs matching the reference. These are conservative analogues of recall and precision: the gold standards are incomplete, so a candidate absent from the reference is not necessarily a false positive. For brevity, table headers and figure axes abbreviate these as *GS recovery* and *GS fraction*. The standard term *recall@20* is retained where it refers to its conventional meaning of ranked-list recovery within the top 20 positions.

**Matching criteria.** A pair "matches" if both entity names match known reference entities, with normalisation for aliases, capitalisation, and common abbreviations.

**Conservative measurement.** All pairs not in the reference set are counted as unmatched. The gold-standard fraction is therefore a lower bound: many unmatched candidates may represent real associations absent from the gold standard, as the plausibility analysis (Section S13) confirms.

**Evaluation views.** Results are reported for (1) all accepted pairs regardless of entity types, and (2) pairs filtered to contain the query entity (e.g., for topic "chondrocyte", only pairs where one entity is "Chondrocyte"). The main text reports the "all accepted" view, which more directly measures gold-standard recovery; the filtered view is reported in per-topic tables below. Each view is in turn reported on two footings: *ungated*, counting every accepted pair, and *on-topic*, counting only pairs the on-topic restriction (Section S1.4) accepts, which is the report's default. Pooled across the 60 topics, the restriction roughly halves the accepted pairs (13,826 to 6,891) while retaining 90.9% of the distinct gold partners recovered (590 of 649), so the default view is markedly smaller at a small recovery cost. Unless stated otherwise, the per-topic, plausibility, and ranking results in the sections below are reported on this on-topic default view.

**On-topic restriction calibration.** Across all 60 case-study topics, the restriction retains 90.9% of gold partners (590 of 649) while cutting 47.5% of off-topic candidate rows (4888 of 10301), scored against the same gold matcher used throughout. The distribution is uneven: most topics retain all of their gold, with a tail of broad-subject and low-gold topics (for example hypertriglyceridemia at 62% and migraine at 75%) sitting lower. In these topics gold is credited to symptoms, mechanisms, genes, or sibling cell-stages, which the restriction cannot admit without also re-admitting the off-topic candidates it exists to remove. We selected GPT-5-mini over the cheaper Haiku because Haiku was markedly stricter on the taxonomy question and lost more gold for the same amount of off-topic removal; a per-pair `topic_relevance` score, though free and already computed, conflates "biologically related" with "is the subject" (it rates smooth muscle cell as highly relevant to neural crest cells), so it judges relevance rather than identity. No single check both retains at least 90% of gold and removes most of the off-topic remainder; the taxonomic subject check is the best option that does not require the model to reason about associations.

**Baseline approaches.** The manuscript compares interaction-finder against two baselines: single-shot direct prompting and DeerFlow (off-the-shelf deep research framework, commit `4459595d`). These were chosen as the simplest and most comparable general-purpose approaches; both were evaluated on the same 20 topics per domain with 5 repeats each. Single-shot prompting used GPT-5-mini; DeerFlow used GPT-4.1-mini due to website rate-limiting constraints. To bound the effect of this model difference, single-shot was additionally run with GPT-4.1-mini: mean gold-standard recovery was 20% vs 28% (celltype–cellmarker), 4% vs 6% (disease–gene), and 29% vs 24% (ligand–receptor) for GPT-4.1-mini vs GPT-5-mini respectively. The model difference is small relative to the gap between baselines and interaction-finder.

Three further approaches (single-shot with provided context, an agentic workflow with information-retrieval tools, and a custom deep-research workflow with specialised prompts) were also evaluated during development and are preserved in the repository, but are not analysed here. The two baselines above are sufficient to establish that interaction-finder's integrated search and extraction substantially outperforms simpler LLM-driven approaches at the end-to-end task; ranking general-purpose LLM strategies against one another is not a goal of this work.

**interaction-finder configuration.** All runs used GPT-5-mini (`openai:gpt-5-mini/low+flex`) as the default model, with GPT-5 (`openai:gpt-5/medium+flex`) for the final cross-document judgment stage. PubMed was the primary search backend; SearXNG was used for keyword-stage review searches. All other settings were left at their default values.

**Variability assessment.** To quantify run-to-run variability, one representative topic per domain was selected and run five times with identical configuration (Table S8). The selected topics (intestinal stem cell, femoral bowing, and ASIP, one per domain) were chosen because their gold-standard fraction and recovery fall near the median for their respective domains, and their reference set sizes span the range encountered across the evaluation (18, 51, and 8 known pairs respectively).

**Table S8:** Run-to-run variability across five runs per topic (full candidate list; this stochasticity is a property of extraction, independent of the on-topic restriction).

| Topic | Run | GS fraction | GS recovery | Cost |
|---|---|---|---|---|
| Intestinal stem cell | 1 | 28.2% | 77.8% | $0.74 |
| | 2 | 24.3% | 77.8% | $11.05 |
| | 3 | 26.5% | 77.8% | $5.78 |
| | 4 | 27.3% | 77.8% | $7.58 |
| | 5 | 27.6% | 66.7% | $4.96 |
| Femoral bowing | 1 | 50.0% | 21.6% | $1.61 |
| | 2 | 45.0% | 45.1% | $3.34 |
| | 3 | 35.4% | 39.2% | $3.67 |
| | 4 | 33.5% | 29.4% | $2.71 |
| | 5 | 29.6% | 17.6% | $1.85 |
| ASIP | 1 | 29.4% | 50.0% | $3.12 |
| | 2 | 52.5% | 50.0% | $1.69 |
| | 3 | 30.3% | 37.5% | $1.79 |
| | 4 | 73.8% | 75.0% | $2.52 |
| | 5 | 61.5% | 75.0% | $2.31 |

The standard deviation of gold-standard recovery across five runs was 5.0% for intestinal stem cell (mean 75.6%, 12 of 18 reference pairs found in all runs, 15 in at least one), 11.6% for femoral bowing (mean 30.6%, 5/51 in all runs, 30 in at least one), and 16.8% for ASIP (mean 57.5%, 3/8 in all runs, 6 in at least one). The higher variability for ASIP and femoral bowing reflects sensitivity to search trajectory when reference sets are small or the topic has sparse literature.

**Search prompts.** Minimal topic descriptions were used, following these templates:

```
1. interaction-finder extract "cell-markers for <cell type>"
   -e cellmarker -e celltype
2. interaction-finder extract "genes associated with <phenotype>"
   -e gene -e phenotype
3. interaction-finder extract "receptors that bind to <ligand>"
   -e receptor -e ligand
```

#### S7 Expanded tool comparison

The manuscript's tool table groups approaches by category with one representative tool per category. The expanded comparison here lists 16 individual tools against six design dimensions, drawing out the specific combinations of choices that distinguish them (Table S9).

Table S9: Expanded comparison of literature mining tools across key dimensions.

| Tool | Entity types | Input | Text scope | Provenance | Infrastructure | Custom types |
|---|---|---|---|---|---|---|
| PubTator 3.0 | 6 fixed | Search indexed | Abstract | Article-level | Web service | No |
| BERN2 | 9 fixed | Document | Abstract | Entity-level | Web service | No |
| DisGeNET | Gene-disease | Database query | Abstract | PMIDs + text | Web service | No |
| DISEASES | Gene-disease | Database query | Abstract | Scores | Web service | No |
| ARROWSMITH | MeSH terms | Interactive | Title | Co-occurrence | Web service | No |
| BITOLA | MeSH terms | Interactive | Abstract | Co-occurrence | Local | No |
| SciLinker | 4 fixed | Batch | Abstract | Sentence | Multi-node | No |
| SKiM-GPT | User terms | Hypotheses | Abstract | RAG chunks | Single machine | Partial |
| ENQUIRE | Gene + MeSH | User corpus | Abstract | Network | Single machine | No |
| SPIRES/OntoGPT | Configurable | Documents | Abstract | Document | Single machine | Yes |
| LLM-IE | Configurable | Documents | Full text | Document | Single machine | Yes |
| SyRACT | Fixed schemas | Documents | Abstract | Document | Single machine | No |
| LORE | Gene-disease | Topic string | Abstract | Statements | Single machine | No |
| FuncFetch | Enzyme-substrate | PubMed query | Full text | Document | Single machine | No |
| BioNExt | BioRED schema | Documents | Abstract | Document | Single machine | No |
| interaction-finder | User-defined | Topic string | Full text | Quote + valid. | Single machine | Yes |

The key differentiators visible in Table S9:

**On-demand vs. document-dependent.** Most extraction tools (SPIRES/OntoGPT, LLM-IE, SyRACT, BioNExt) require documents as input; they perform extraction but not discovery. LORE and interaction-finder both accept topic strings, but LORE is restricted to disease–gene associations from abstracts. FuncFetch accepts PubMed queries but is domain-specific (enzyme families) without iterative search refinement.

**Quote-level provenance.** Among the tools compared here, interaction-finder is unique in validating extracted quotes against source documents. DisGeNET provides representative sentences, SciLinker identifies co-occurrence sentences, and SPIRES links to ontology terms, but none verify that LLM-extracted content faithfully represents the source text.

**Custom entity types.** SPIRES/OntoGPT, LLM-IE, and interaction-finder support user-defined entity types. However, SPIRES and LLM-IE require documents and do not include literature discovery.

Supervised RE systems like BioREX (Lai et al. 2023) achieve 79.6% F1 on BioRED (Islamaj et al. 2024), setting the performance ceiling for fixed-schema extraction on curated corpora. However, supervised approaches require annotated training data that does not exist for arbitrary entity types: the fundamental trade-off motivating interaction-finder's zero-shot approach.

The combined search-then-extract design places interaction-finder in the emerging class of agentic deep research systems (Gridach et al. 2025; Huang et al. 2025). FuncFetch (Smith et al. 2025) demonstrates that this pattern can work in biological domains, achieving 0.86/0.64 precision/recall on enzyme-substrate extraction from full-text manuscripts, though for a single relationship type. The Literature Search Sandbox (Adam et al. 2024) quantifies the limitation of static LLM-generated queries for systematic review, achieving median 85% sensitivity but low precision (NNR ~1206). The iterative search with LLM reflection in interaction-finder addresses this by adaptively expanding coverage based on gap assessment between rounds.

### S8 Per-topic evaluation tables

#### S8.1 Celltype–cellmarker

Table S10: Per-topic results for celltype–cellmarker domain (all accepted pairs view).

| Topic | Ref. pairs | GS fraction | GS recovery | Cost |
|---|---|---|---|---|
| Activated B cell | 17 | 17.4% | 76.5% | $5.62 |
| Alveolar type II cell | 9 | 12.9% | 66.7% | $0.78 |
| Capillary cell | 14 | 9.3% | 57.1% | $3.86 |
| Chondrocyte | 7 | 10.3% | 85.7% | $11.61 |
| Common lymphoid progenitor cell | 6 | 16.4% | 66.7% | $2.50 |
| Dopaminergic neuron | 11 | 18.5% | 54.5% | $2.57 |
| Effector T cell | 18 | 25.0% | 61.1% | $5.85 |
| Endocardial cell | 12 | 13.6% | 83.3% | $3.38 |
| Erythroblast | 9 | 13.8% | 66.7% | $4.79 |
| Hepatoblast | 11 | 18.5% | 90.9% | $4.31 |
| Intestinal stem cell | 18 | 24.1% | 77.8% | $0.74 |
| Intestinal stem cell (GPT-5) | 18 | 31.7% | 72.2% | $63.16 |
| Intestinal stem cell (GPT-5-nano) | 18 | 43.2% | 44.4% | $1.77 |
| Kupffer cell | 12 | 15.0% | 58.3% | $3.35 |
| Langerhans cell | 13 | 26.1% | 69.2% | $6.16 |
| Limbal stem cell | 12 | 11.9% | 75.0% | $5.66 |
| Mature luminal cell | 6 | 18.6% | 50.0% | $2.99 |
| Mature neutrophil | 11 | 18.5% | 45.5% | $2.16 |
| Megakaryocyte progenitor cell | 5 | 7.2% | 60.0% | $4.54 |
| Neural crest cell | 35 | 20.4% | 48.6% | $14.21 |
| Plasmablast | 28 | 25.8% | 53.6% | $3.96 |
| Proximal tubular cell | 20 | 12.9% | 60.0% | $3.94 |

Per-topic celltype–cellmarker results are shown in Table S10. The three intestinal stem cell runs illustrate cost-quality tradeoffs: GPT-5-mini achieves the highest gold-standard recovery (77.8%) at the lowest cost ($0.74), GPT-5 provides a modest gold-standard fraction improvement (31.7% vs 24.1%) at 85× the cost ($63.16), and GPT-5-nano improves gold-standard fraction (43.2%) but at the expense of recovery (44.4%) and quote fidelity (86.9% validation rate). That model choice substantially affects biomedical NLP performance is well established (Chen et al. 2025), and interaction-finder's hierarchical model configuration (Section S11) lets users balance cost against quality per pipeline stage.

Gold-standard recovery ranges from 45.5% (mature neutrophil) to 90.9% (hepatoblast), with a mean of 65.3% across the 20 primary topics. The gold-standard fraction ranges from 7.2% (megakaryocyte progenitor cell) to 26.1% (Langerhans cell). Per-topic cost ranges from $0.74 to $14.21, with the variation largely reflecting the volume of discoverable literature for each cell type.

**Table S11:** Per-topic results for disease–gene domain (all accepted pairs view).

| Topic | Ref. pairs | GS fraction | GS recovery | Cost |
|---|---|---|---|---|
| Autoimmune thrombocytopenia | 48 | 14.0% | 27.1% | $2.65 |
| Dilated cardiomyopathy | 185 | 41.1% | 29.2% | $11.49 |
| Emphysema | 63 | 11.1% | 20.6% | $14.21 |
| Episodic paroxysmal anxiety | 17 | 0.0% | 0.0% | $3.15 |
| Femoral bowing | 51 | 51.7% | 15.7% | $1.61 |
| Hypercholesterolemia | 85 | 55.4% | 12.9% | $6.63 |
| Hypertriglyceridemia | 133 | 59.2% | 9.8% | $4.99 |
| Migraine | 155 | 18.1% | 9.7% | $7.20 |
| Nephrotic syndrome | 137 | 60.2% | 35.8% | $11.93 |
| Parkinsonism | 141 | 39.2% | 24.1% | $8.38 |
| Pituitary hypothyroidism | 38 | 45.5% | 44.7% | $0.61 |
| Progressive hearing impairment | 60 | 23.9% | 30.0% | $8.30 |
| Psoriasiform dermatitis | 33 | 9.4% | 21.2% | $5.88 |
| Pulmonary arterial hypertension | 164 | 12.8% | 26.8% | $1.52 |
| Pulmonary fibrosis | 78 | 24.2% | 34.6% | $7.75 |
| Renal tubular acidosis | 49 | 35.0% | 20.4% | $3.25 |
| Rhabdomyolysis | 45 | 46.9% | 51.1% | $6.75 |
| Rheumatoid arthritis | 23 | 5.5% | 47.8% | $12.58 |
| Truncal ataxia | 77 | 33.3% | 14.3% | $3.68 |
| Zonular cataract | 52 | 35.2% | 40.4% | $0.25 |

#### S8.2 Disease–gene

Per-topic disease–gene results are shown in Table S11. Gold-standard recovery falls as reference set size grows: topics with small reference sets (≤52 genes) recover up to around half of their pairs, while the four largest (≥141 genes) recover only 10–29%. On-the-fly literature discovery cannot keep pace once reference sets reach into the hundreds, regardless of the underlying biology.

Episodic paroxysmal anxiety is the sole topic with 0% gold-standard recovery, and illustrates a gold-standard mismatch: interaction-finder extracted 198 accepted pairs predominantly involving anxiety-related genes (e.g. BDNF, SLC6A4, NPSR1, SCN1A), but the HPO reference maps this phenotype to pheochromocytoma and paraganglioma genes (VHL, RET, SDHB, SDHD, NF1) via the ORPHA disease ontology. The reference association is indirect: pheochromocytoma can present as episodic anxiety attacks, and the genes that the literature directly associates with anxiety disorders are entirely absent from the reference set. The 0% therefore reflects a mismatch between the reference standard's ontological mapping and the literature's direct gene–phenotype associations, more than a failure of the extraction pipeline.

The gold-standard fraction in the disease–gene domain is higher than in celltype–cellmarker (mean 31% vs 17%); we return to the cross-domain comparison after presenting ligand–receptor.

#### S8.3 Ligand–receptor

**Table S12:** Per-topic results for ligand–receptor domain (all accepted pairs view).

| Topic | Ref. pairs | GS fraction | GS recovery | Cost |
|---|---|---|---|---|
| ADAM2 | 5 | 38.5% | 80.0% | $0.74 |
| AREGB | 2 | 0.0% | 0.0% | $2.01 |
| ASIP | 8 | 33.3% | 25.0% | $3.12 |
| CGA | 4 | 6.7% | 25.0% | $1.75 |
| CSF3 | 2 | 100.0% | 50.0% | $1.08 |
| CXCL6 | 2 | 80.0% | 100.0% | $1.57 |
| JAG2 | 4 | 75.0% | 50.0% | $1.99 |
| MFAP2 | 1 | 25.0% | 100.0% | $0.92 |
| OBP2A | 1 | 0.0% | 0.0% | $2.12 |
| PDAP1 | 1 | 0.0% | 0.0% | $1.59 |
| PROS1 | 2 | 33.3% | 100.0% | $1.94 |
| PSEN1 | 6 | 0.0% | 0.0% | $2.32 |
| PTHLH | 3 | 46.7% | 66.7% | $4.03 |
| RAET1E | 1 | 0.0% | 0.0% | $0.96 |
| SEMA6D | 4 | 22.2% | 75.0% | $1.14 |
| SERPINA7 | 1 | 0.0% | 0.0% | $1.34 |
| SFTPD | 2 | 0.0% | 0.0% | $0.68 |
| SLPI | 1 | 0.0% | 0.0% | $0.63 |
| TSHB | 1 | 60.0% | 100.0% | $4.76 |
| VEGFB | 4 | 37.5% | 50.0% | $4.04 |

Per-topic ligand–receptor results are shown in Table S12. The mean gold-standard recovery across all 20 topics is 41.1%, with 12 of 20 ligands recovering at least one known receptor. Four ligands achieve 100% recovery (CXCL6, MFAP2, PROS1, TSHB, all with 1–2 known receptors), while eight ligands with sparse or unfindable literature, or whose accepted pairs the restriction marks off-topic, recover 0% (OBP2A, RAET1E, SERPINA7, SLPI, SFTPD, AREGB, PDAP1, PSEN1). The on-topic restriction empties three of these topics of candidates entirely; the main-text's Table 2 reports the mean over the 17 topics that retain candidates (48%), so it sits above this all-20 average. The gold-standard fraction in this domain (mean 28%) remains close to disease–gene (31%) and exceeds celltype–cellmarker (17%). In both disease–gene and ligand–receptor, the reference relationship typically names specific molecular partners (a gene of a disease, a receptor of a ligand) that appear explicitly in the literature; cellmarker references include broadly expressed proteins that the literature mentions in many contexts, inflating the count of accepted pairs that fall outside the reference and depressing the gold-standard fraction.

#### S8.4 Cross-topic analysis

**Score non-discrimination.** Across all 60 evaluated topics, LLM-assigned evidence scores and confidence scores show nearly identical distributions for gold-standard matches and unmatched candidates (Fig. S2). Among the representative topics the largest difference (VEGFB: matched mean 7.67 vs unmatched 7.04) remains well within one standard deviation, and in several topics unmatched candidates score marginally higher than matched (for example PTHLH, 7.10 vs 7.22). The same pattern holds for confidence scores (not shown). Document count (see Fig. S3) provides a more effective ranking signal, considered next.

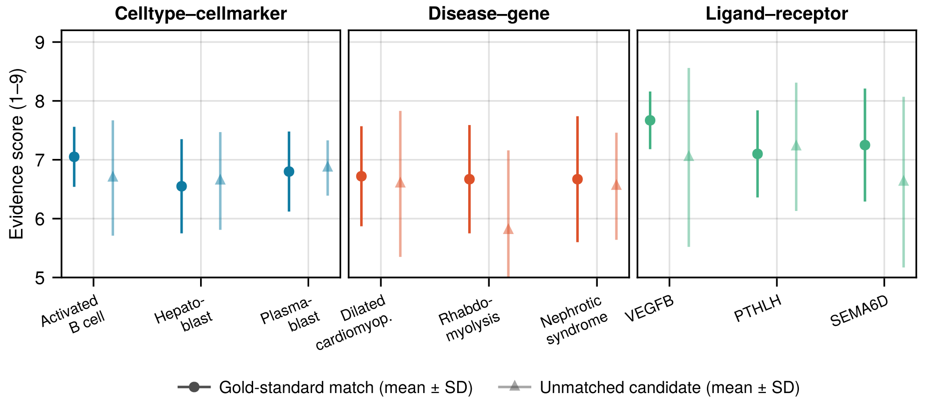

**Figure S2:** Evidence score distributions for gold-standard matches (circles) and unmatched candidates (triangles) across representative topics from each domain (mean ± standard deviation).

**Document count as ranking signal.** Filtering by minimum document count consistently improves the gold-standard fraction at the expense of recovery (Fig. S3). The tradeoff is steepest in the disease–gene domain, where aggressive filtering (≥10 documents) roughly doubles the gold-standard fraction but retains only a small portion of known associations. For celltype–cellmarker and ligand–receptor topics, moderate filtering (≥3–5 documents) yields a more favourable tradeoff. The gold-standard fraction here treats every unmatched candidate as a false positive; the plausibility analysis (Section S13) suggests true precision is substantially higher across all thresholds.

The pattern is consistent across all three domains: higher document-count thresholds improve the gold-standard fraction at the expense of recovery, with the tradeoff varying by domain. For celltype–cellmarker and ligand–receptor topics, the recovery cost is moderate, making document count a practical ranking signal for expert triage. In the disease–gene domain, where reference sets are much larger, the recovery cost of aggressive filtering is more severe (filtering to ≥10 documents typically retains only 5–20% of known associations), reflecting the fact that many disease–gene relationships appear in only a few papers.

**Keyword-stage enrichment.** Because the keyword-discovery stage mines review articles, which tend to mention well-established associations, pairs first discovered during this stage have a consistently higher gold-standard fraction than those found in later search rounds (Table S13). In the disease–gene domain, the keyword-stage fraction (37.6%) is well above that of the first ten wide-search rounds (21.9%); the same holds for celltype–cellmarker (24.1% vs 15.2%) and ligand–receptor (32.9% vs 24.4%). The fraction generally declines in later rounds as the pipeline reaches into more peripheral literature. The wide variance in the later rounds (e.g. 0–100% for ligand–receptor) reflects the diminishing

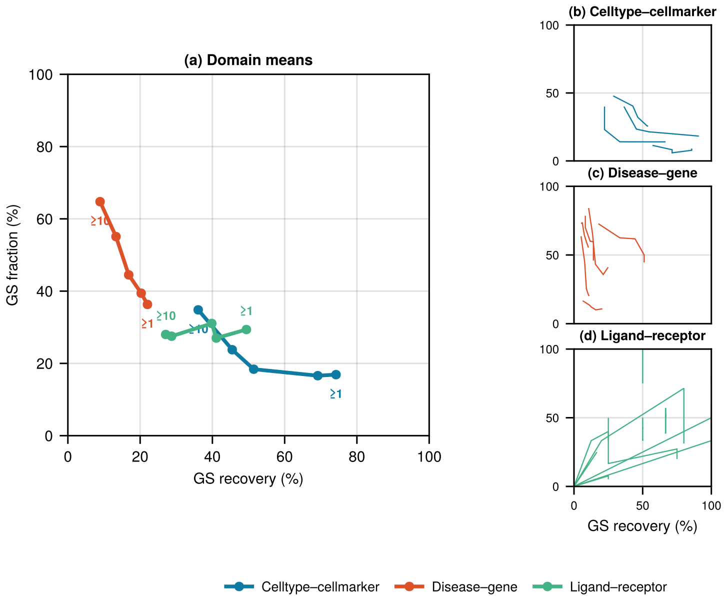

**Figure S3:** Gold-standard fraction and recovery under document-count filtering. **(a)** Mean gold-standard fraction and recovery across 20 topics at each threshold (≥1 through ≥10). **(b)**–**(d)** Per-topic curves for celltype–cellmarker, disease–gene, and ligand–receptor respectively, with one line per topic.

and increasingly variable returns from extended searching; for ligand–receptor in particular the later rounds retain only a handful of pairs per bin, so their per-round fractions are dominated by sampling noise rather than a stable trend.

**Table S13:** Gold-standard fraction by search round provenance (combined across all topics per domain). Keyword-stage pairs originate from review articles mined during the initial keyword discovery phase; subsequent rounds correspond to the iterative wide search. Each round column groups 10 consecutive search queries; the "Later rounds" column reports the range across per-domain late-query bins (queries 21+, in further groups of 10), reflecting the diminishing volume and increasing variability of pairs first discovered at these depths.

| Domain | Keywords | Rounds 1–10 | Rounds 11–20 | Later rounds |
|---|---|---|---|---|
| Celltype-cellmarker | 24.1% | 15.2% | 12.2% | 9–17% |
| Disease-gene | 37.6% | 21.9% | 19.4% | 4–25% |
| Ligand-receptor | 32.9% | 24.4% | 36.4% | 0–100% |

#### S9 Default sort in the interactive report

The report needs a sensible default order for candidates. Document count is a natural choice, but it privileges pairs with long literature trails: the pairs most likely to already appear in curated databases, and therefore the pairs an interaction-finder user is least likely to be looking for. The curated gold standards used here (Ramilowski et al. 2015; Hu et al. 2023; Gargano et al. 2024) inherit this bias. Any ranker that accounts for novelty is penalised by gold-only evaluation because it spends ranking positions on pairs the evaluation treats as false positives; the magnitude of this bias is large (Mayers et al. (2019) report cross-validation AUROC of 0.95 on their network drug-repurposing model dropping to 0.797 when restricted to post-cutoff novel associations), and the literature-based-discovery community has converged on temporal holdout evaluation to address it (Thilakaratne, Falkner, and Atapattu 2019; Moreau 2023). We evaluated candidate sort rules on two complementary axes: recall@20 against the full gold standard, and recall@20 against the subset of gold pairs whose earliest supporting document in the pipeline's retrieved corpus is from 2018 or later, as a proxy for novel discovery. The proxy is calibrated to what the pipeline saw rather than true first-appearance in literature, but is fair for comparative purposes because the calibration is identical across rankers.

##### S9.1 Evaluation protocol

Recall@20 is computed per topic and averaged across topics (macro-average). Splits are topic-stratified at 70/30 (42 training, 18 test topics per split), with topics assigned uniformly at random subject to each of the three domains contributing train and test topics. The 70/30 split was resampled under 50 seeds for bootstrap estimates. Topic-stratification rather than row-stratification is essential: pairs within a topic are not independent, so row-stratified splits leak the topic's signal between train and test and inflate every ranker equivalently, hiding their differences.

##### S9.2 Candidate rules, and why most combinations failed

We tested four ranker families: single-feature sorts (document count, the judge's pair-level topic relevance, the maximum per-document specificity, a recency-weighted sum of supporting documents), linear combinations fitted to gold membership by logistic regression, gradient-boosted ensembles over 67 per-pair features, and rank sums combining strong single features. A rank sum of two features (summing each feature's within-topic rank) outperformed every other family on held-out topics.

Linear combinations of the two best single features collapsed into a one-feature regime under every combination rule we tested. The cause is distributional: the z-scored recency-weighted document sum has typical within-topic range [−1.3, +13.7] because the underlying sum is heavy-tailed, while other relevance features are bounded integers with most pairs clustered near the maximum. Any linear combination is dominated by the feature with larger effective range. Summing within-topic ranks sidesteps the issue by putting both features on the same bounded scale before combining.

The gradient-boosted model offered no advantage for surfacing curated pairs: trained against the independent-judge P(TP) target (Section S13) it reached held-out Spearman 0.48 against that target, but produced an R@20 of 0.60 on full gold, no better than the simple single-feature sorts despite its far greater complexity and its dependence on a pretrained model. Predicting a continuous plausibility score and surfacing pairs present in the curated reference are genuinely different objectives.

#### S9.3 The chosen ranking rule

Each candidate pair is scored as the sum of two within-topic ranks: one over a *quote-weighted relevance* score, and one over a *recency-weighted document* score. We define each in turn before giving the combined formula.

**Quote-weighted relevance.** Each pair has one per-document assessment $i$ for every document that mentions both entities. The assessment carries a topic-relevance rating $t_i \in \{1, \ldots, 5\}$ and supporting quotes of total character length $L_i$. Substantiation is encoded by combining the two into

$$s_i = t_i \cdot \left(1 - e^{-L_i/200}\right).$$

The pair's relevance feature $R_{\text{pair}}$ is the mean of the raw ratings $t_i$ for the three assessments with the largest $s_i$ (i.e. $s_i$ is used only to pick the top three; the returned value is the average of their raw ratings, not their substantiated scores).

**Recency-weighted document sum.** Let $D$ be the set of unique documents supporting the pair, and $\text{age}(d)$ the age of document $d$ in years between its publication and report generation. The recency feature is

$$\text{rec}_{\text{pair}} = \sum_{d \in D} (\text{age}(d) + 1)^{-\frac{3}{4}}.$$

**Combined score.** The candidate's overall score is the sum of the within-topic ranks of these two features:

$$\text{score(pair)} = \text{rank}_{\text{topic}}(R_{\text{pair}}) + \text{rank}_{\text{topic}}(\text{rec}_{\text{pair}}).$$

Ranks are assigned within each topic in descending order of the underlying value (ties get the average rank), and the pair with the lowest total rank is presented first. The recency-weighted document sum alone gives the best recall on recent pairs (see Table S15), so the report defaults to that single feature; the two-feature rank-sum above is retained as a selectable alternative that weights substantiated relevance equally with recency. The substantiation weight $1 - \exp(-L/200)$ was selected from a grid over exponential, Hill, logistic, and soft-threshold forms; the characteristic length $\tau = 200$ characters matches the binned-means peak of gold-membership fraction against quote length (peak at 150–200 chars, see Section S9.4 below). Alternative $\tau$ values in $[100, 300]$ give equivalent performance.

The recency feature sums $(\text{age} + 1)^{-3/4}$ across unique supporting documents, with age in years between the document's publication and report generation. The exponent 3/4 was selected by sweeping the inverse-power decay family $(1/(\text{age} + 1)^p)$ under topic-stratified cross-validation: it improves held-out $\geq 2020$ recall@20 over the milder $1/\sqrt{\text{age} + 1}$ decay we initially used (+0.10, winning 92% of folds) at no cost to full-gold recall (see Section S9.5 below). We retained an inverse-power form, rather than exponential decay ($\exp(-\text{age}/\tau)$ for $\tau \in [1, 50]$, which gave equivalent performance across its optimal range), for its long tail, which lets older papers continue to contribute meaningfully.

#### S9.4 Quote-length signal

The substantiation weight is motivated by the binned-means relationship between mean supporting-quote length and each evaluation axis, across accepted pairs (Table S14). Shorter quotes are less substantive; the peak substantiation occurs around 150–200 characters, after which longer quotes do not further increase the gold-membership rate or mean plausibility. A saturating weight function matches this shape; an unbounded monotone function (e.g. $\sqrt{L}$) overweights long quotes relative to the empirical peak.

**Table S14:** Binned metric means by mean supporting-quote length, pooled across accepted pairs. `frac_gold` is the fraction of pairs in the gold reference. `frac_novel_2020` is the fraction of pairs whose earliest supporting document is from 2020 or later. `mean_plaus` is the mean plausibility rating on the 1–9 scale (Section S13).

| Bucket (chars) | n | frac_gold | frac_novel_2020 | mean_plaus |
|---:|---:|---:|---:|---:|
| 1–50 | 21 | 0.048 | 0.048 | 3.62 |
| 51–100 | 276 | 0.091 | 0.047 | 4.60 |
| 101–150 | 1229 | 0.104 | 0.038 | 4.96 |
| 151–200 | 1429 | 0.116 | 0.052 | 5.28 |
| 201–300 | 590 | 0.058 | 0.039 | 4.89 |
| 301–500 | 95 | 0.105 | 0.084 | 4.90 |

#### S9.5 Results

**Table S15:** Recall@20 for candidate default sort rules, averaged across 50 topic-stratified 70/30 bootstrap splits (mean ± standard deviation). The "≥2020" column evaluates against gold pairs whose earliest supporting document in the pipeline corpus is from 2020 or later. The recency-weighted document sum, which gives the best recall on recent (≥2020) pairs, is the interactive report's default.

| Sort rule | Full gold | ≥2020 |
|---|---|---|
| Document count | 0.626 ± 0.079 | 0.412 ± 0.132 |
| Pair-level topic relevance | 0.604 ± 0.079 | 0.420 ± 0.122 |
| **Recency-weighted doc sum** | **0.610 ± 0.077** | **0.569 ± 0.122** |
| Rank-sum of $R_{\text{pair}}$ + recency | 0.620 ± 0.075 | 0.491 ± 0.111 |

Averaged across 50 bootstrap splits, the four rules are tightly clustered on the full reference (0.604–0.626 R@20, all within one bootstrap standard deviation of each other), with document count nominally highest. The rules separate on the ≥2020 novel subset, where recency-aware ranking helps: the recency-weighted document sum leads at 0.569 ± 0.122, well ahead of the rank-sum fusion (0.491) and document count (0.412). The marginal standard deviations are large because they are dominated by topic-to-topic difficulty; the relevant test is the paired difference across matched splits. The $(\text{age}+1)^{-3/4}$ decay was chosen over the milder $1/\sqrt{\text{age}+1}$ we initially used, which underweights recent work. To guard against tuning on the evaluation set, we confirmed the choice under topic-stratified cross-validation (select the exponent on training topics, evaluate on held-out topics, 50 splits): against the $1/\sqrt{\text{age}+1}$ baseline the steeper decay improves held-out ≥2020 recall@20 by +0.10 (0.44 to 0.54), winning 92% of folds, at no cost to full-gold recall (+0.006, indistinguishable from zero). The gain rests on a modest 171 recent gold pairs, so the per-fold variation is wide, but the direction is consistent across folds. The recency-weighted sum is therefore the default of choice for a discovery tool: a consistent gain on the recent pairs that matter most while remaining a simple, interpretable single-feature rule.

A larger Pareto sweep across 17 244 alternative weighted rank-sum combinations of uncorrelated feature pairs and triples (using features such as spec_max, evidence, ndocs, paper quality) found no combination that strictly dominated the recommended rule on all four evaluation axes (full gold, ≥2015 novel, ≥2020 novel, top-20 mean plausibility) on the ungated candidate set. Further sweeps varying the per-document topic-relevance aggregation (power sums, threshold sums, top-K means, quantile-based) and the substantiation weight function (exponential, logistic, Hill, soft-threshold) produced no combination that dominated the $\tau = 200$ exponential with top-3 mean aggregation on more than one axis while maintaining performance on the others. Once off-topic candidates are removed the rule ordering tightens and the rank-sum fusion no longer holds a clear edge: the recency-weighted document sum gives the best recall on the recency-restricted axes (≥2020 novel recall@20 of 0.569, versus 0.491 for the rank-sum fusion), and it is the report's default for that reason, most directly surfacing the recent candidates that matter for discovery while remaining a simple, single-signal rule. Five formulations that fuse recency into the per-assessment score (rather than ranking it separately) underperform the separate-features form, because fusing recency at the assessment level mutes the recency signal for pairs with a few old well-substantiated assessments that would otherwise dominate the selection.

The recency-weighted default trades a small loss in recall of long-established associations for earlier visibility of recent candidates. Users focused on established associations can re-sort by document count; the interactive report exposes both orderings without re-generating the report.

#### S9.6 Per-rank enrichment of plausibility and gold-standard pairs

Recall@20 reduces the ranking to a single number per topic. Fig. S4 and Fig. S5 show the default recency-weighted ranker against two complementary signals across the full rank range.

Fig. S4 shows the mean plausibility rating at each rank, computed twice per domain: once over all candidates including gold-standard recapitulations, and once over the non-gold subset only. Both curves drop from approximately 7 at the top of the list to the topic baseline by rank 50–100 across disease–gene and celltype–cellmarker, and remain elevated through the full rank range for ligand–receptor (which has shorter candidate lists per topic). The two curves remain close in level and shape throughout the rank range, so non-gold candidates near the top of the list are scored just as plausibly as the gold-standard recapitulations there; the falling curve reflects a property of the candidate population as a whole, not an artefact of gold-standard pairs being preferentially surfaced.

Fig. S5 shows the per-rank gold-standard hit rate alongside the rate that would obtain under a post-hoc ranking by Method-B plausibility (an oracle that cannot be computed without the independent judge). The recency-weighted default ranker tracks the plausibility-ranked oracle closely across all three domains, indicating that it captures most of the signal a plausibility-aware ranker could provide while using only features available at extraction time. The uniform baseline accounts for the changing set of contributing topics at each rank, so the gap between the ranker's curve and the baseline is a fair measure of enrichment.

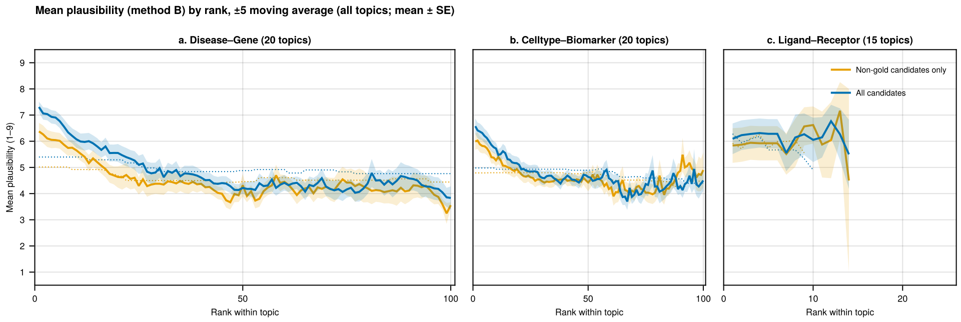

**Figure S4:** Mean plausibility rating (Method B, 1–9 scale) of accepted candidates against their rank in the default sort, smoothed with a ±5 rank window, for **(a)** disease–gene, **(b)** celltype–cellmarker, and **(c)** ligand–receptor. The blue trace pools all candidates including gold-standard recapitulations; the orange trace covers non-gold candidates only. The dotted lines are rank-conditional baselines, averaging each subset's topic-mean rating over only the topics still contributing at each rank (matching the baseline treatment in Fig. S5).

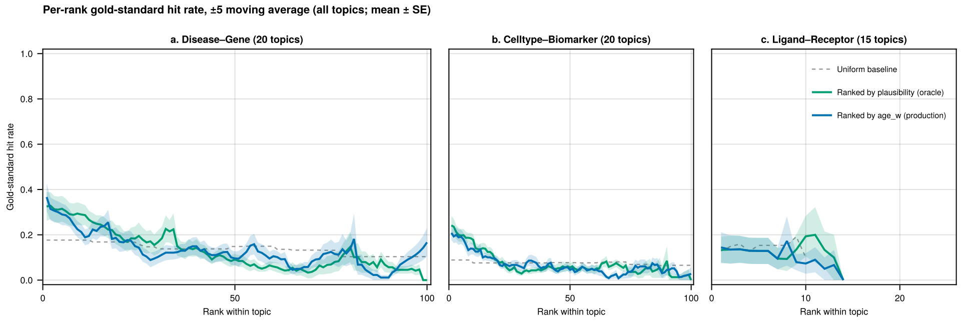

**Figure S5:** Per-rank gold-standard hit rate (±5 rank moving average) for **(a)** disease–gene, **(b)** celltype–cellmarker, and **(c)** ligand–receptor, comparing the rank-sum ranker (blue), a post-hoc ranking by Method-B plausibility (green), and uniform random ranking (grey dashed). The uniform baseline averages over only the topics still contributing at each rank.

#### S10 Contamination experiment

##### S10.1 Design and document sampling

For each of the 60 evaluated topics, 20 documents were sampled from a different domain's completed search results and injected into the target topic's corpus. The cross-domain mapping (Table S16) was chosen so that source documents contain real biological content but of the wrong kind for the target topic (e.g., ligand–receptor articles spiked into disease–gene topics).

**Table S16:** Cross-domain spiking scheme. Pool size is the number of deduplicated documents with non-null content available for sampling in each source domain.

| Target domain | Target entity types | Source domain (spiked) | Pool size |
|---|---|---|---|
| Disease–gene | gene, phenotype | Ligand–receptor | 2,171 |
| Celltype–cellmarker | cellmarker, celltype | Disease–gene | 5,223 |
| Ligand–receptor | ligand, receptor | Celltype–cellmarker | 3,064 |

Source pools were constructed by collecting all resources with non-null content from the base-run checkpoints of each source domain, deduplicating by URL. For each target topic, 20 documents were sampled uniformly at random using a deterministic per-topic seed (`hash("42-{topic}") mod 2^{32}`), after excluding any URL already present in the target checkpoint. The resulting contamination fraction ranged from 3.4% (pulmonary arterial hypertension, 577 real documents) to 33.9% (ADAM2, 39 real documents), with a mean of 13.8% across all 60 topics.

##### S10.2 Per-domain extraction funnel

Table S17 breaks down the extraction funnel by domain. Entity extraction is the dominant filter in all three domains: 92–96% of spiked documents produce no entities of the target type, against 31.5% of real documents (68.5% yield, table). Among spiked documents that do yield entities, roughly half produce assessed pairs. The pipeline's specificity is therefore acting on irrelevant content, not penalising real documents through general stringency. Per-document intermediate stages (proximal sets, extracted pairs) are not tracked in the original checkpoints, so the real-document comparison is limited to entity yield and assessment rates.

Real-document comparison is therefore limited to the entity-yield and assessment-rate stages, since per-document intermediate stages were not retained in the original checkpoint format.

##### S10.3 Per-topic breakdown

Fig. S6 shows, for each of the 60 topics, the number of spiked documents that yielded any target-type entities (light bars) and the number of accepted pairs that resulted (solid bars). Of the 60 topics,

**Table S17:** Per-document extraction funnel for spiked documents, broken down by domain. Entity extraction filters 92–96% of spiked documents before pair assessment, against 31.5% for real documents (i.e. 68.5% of real documents yield target-type entities, last column). The "—" entries in the Real column mark per-document intermediate stages that were not retained in the original checkpoint format. CC = celltype–cellmarker, DG = disease–gene, LR = ligand–receptor.

| Stage | CC (400 docs) | DG (400 docs) | LR (400 docs) | All spiked (1,200) | Real (10,550) |
|---|---|---|---|---|---|
| With target-type entities | 31 (7.8%) | 31 (7.8%) | 16 (4.0%) | 78 (6.5%) | 7,224 (68.5%) |
| With proximal sets | 24 (6.0%) | 25 (6.2%) | 11 (2.8%) | 60 (5.0%) | — |
| With extracted pairs | 17 (4.2%) | 13 (3.2%) | 10 (2.5%) | 40 (3.3%) | — |
| With assessed pairs | 17 (4.2%) | 13 (3.2%) | 10 (2.5%) | 40 (3.3%) | 7,224 (68.5%) |
| Total assessments | 47 | 26 | 38 | 111 | 76,316 |
| Assessments per doc | 0.12 | 0.07 | 0.10 | 0.09 | 7.23 |
| Mean evidence score (1–9) | 6.5 | 6.4 | 6.6 | 6.5 | 6.7 |
| Mean topic relevance (1–5) | 4.1 | 4.1 | 4.4 | 4.2 | 4.6 |

37 produced no spiked entities at all, and most of the remainder produce only one or two accepted pairs. Almost all of the accepted-pair mass sits in a handful of hotspots (endocardial cell, PSEN1, and CGA) where the source domain incidentally discusses entities relevant to the target domain (endothelial markers in vascular-disease papers, Notch ligand–receptor pairs in hepatoblastoma reviews; see Section S10.4 for the per-paper breakdown). The contamination signal is therefore not a pipeline-wide leak but a localised consequence of cross-domain biology overlap, and even at the hotspots the accepted pairs are biologically valid relationships rather than fabrications.

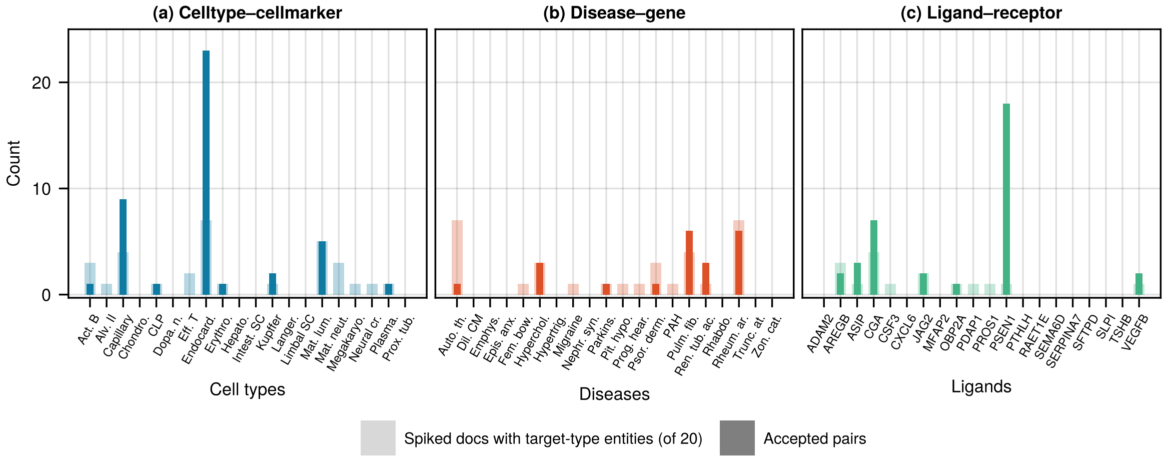

**Figure S6:** Per-topic contamination results for **(a)** celltype–cellmarker, **(b)** disease–gene, and **(c)** ligand–receptor. Light bars show the number of spiked documents (out of 20) that produced target-type entities; solid bars show the number of accepted pairs.

#### S10.4 Source papers producing accepted pairs

Table S18 summarises the 21 source papers that produced at least one accepted spiked pair. The extracted relationships span the full range from off-topic to directly relevant: a B cell paper injected into the ASIP ligand–receptor topic yielded three BAFF receptor pairs unrelated to ASIP, while a proximal tubule endocytosis review injected into the renal tubular acidosis topic yielded three known RTA genes (OCRL, CTNS, CLCN5) not found by the original search. Two papers dominate the count: a hepatoblastoma review accounts for 18 pairs (Notch ligand–receptor interactions, spiked into PSEN1) and a cerebral small vessel disease review accounts for 12 (endothelial and glial markers, spiked into endocardial-cell). In every case the extracted relationships are biologically correct; they arise because cross-domain papers incidentally discuss entities relevant to the target domain.

#### S10.5 Gold-standard matching and outcome stability

Of the 99 accepted spiked-only pairs, 29 (29%) match gold-standard reference datasets under the same evaluation criteria used for the main gold-standard analysis (4 in celltype–cellmarker, 5 in disease–gene, 20 in ligand–receptor). Because spiked entities were not consolidated with real-document entities, this comparison relies on exact name matching and is a lower bound. Most of these 29 matches correspond to associations already found under different entity names in the original results; the exceptions are two Notch pathway pairs in the PSEN1 topic where the original run used generic names ("Jagged", "Notch receptor") while the spiked paper provided specific names (JAG1, DLL4) that matched the reference dataset.

Applying the on-topic restriction to the 99 accepted spiked-only pairs collapses 70 of them (71%) as off-topic, retaining 29: 18 of 43 in celltype–cellmarker, 9 of 21 in disease–gene, and 2 of 35 in ligand–receptor. The restriction is most aggressive in ligand–receptor, where the spiked pairs are furthest from the topic's ligand subject. This is a display-time filter on top of the entity-typing specificity reported above, and it removes the majority of contamination that reaches the accept stage; the pairs it retains are those whose subject-side entity genuinely denotes the spiked topic (for example, the PSEN1 Notch-pathway pairs above). Note these counts are distinct from the gold-match counts: a contaminated pair can match a gold relation from its original domain yet still be off-topic for the spiked topic, and the restriction correctly drops such pairs.

The full set of 111 spiked-pair assessments (main text) decomposes as 99 spiked-only pairs accepted, 9 spiked-only pairs rejected, and 3 mixed-source pairs that combined evidence from both spiked and real documents. In all three mixed-source cases the accept/reject outcome was unchanged relative to the original un-spiked run; no originally rejected pair was flipped to accepted by spiked evidence. The nine rejected spiked pairs all carried weak evidence ($\leq$4/9) and generic relationship labels (all `associated_with`).

**Table S18:** Source papers contributing accepted spiked pairs, grouped by target domain. In every case the extracted relationships are biologically correct; they arise because cross-domain papers incidentally discuss entities relevant to the target domain. "Pairs" counts accepted pairs only. CC = celltype–cellmarker, DG = disease–gene, LR = ligand–receptor. Eight additional single-pair papers are omitted for brevity.

| Target topic | Source paper | From | Pairs | Cross-domain link |
|---|---|---|---|---|
| Capillary cell | Molecular Changes Implicate Angiogenesis and Arterial Remodeling in Systemic Sclerosis-Associated and Idiopathic Pulmonary Hypertension | PAH | 5 | Capillary endothelial markers (CA4, FCN3) in pulmonary disease |
| " | Airway administration of vascular endothelial growth factor siRNAs induces transient airspace enlargement in mice | emphysema | 3 | VEGF and apoptosis markers in alveolar septal cells |
| Endocardial cell | Cerebral small vessel disease: Pathological mechanisms and potential therapeutic targets | migraine | 12 | Endothelial markers (claudin-5, occludin, eNOS) and glial responses |
| " | Deficiency of cold-inducible RNA-binding protein exacerbated monocrotaline-induced pulmonary artery hypertension through Caveolin1 and CAVIN1 | PAH | 3 | Pulmonary endothelial/smooth muscle markers (CD31, SMA) |
| " | Bone morphogenetic protein receptor II is a novel mediator of endothelial nitric-oxide synthase activation | PAH | 2 | eNOS and caveolin-1 in pulmonary artery endothelial cells |
| " | Targeted inhibition of disheveled PDZ domain via NSC668036 depresses fibrotic process | pulm. fibrosis | 2 | $\alpha$-SMA in myofibroblasts and fibroblasts |
| " | Enhanced detection of expanded repeat mRNA foci with hybridization chain reaction | truncal ataxia | 2 | Flag/GFP in mouse embryonic fibroblasts (experimental markers) |
| " | Lymphatic incorporated biomimetic scaffold enhances Osteoangiolymphogenic coupling via HIF-1$\alpha$ mediated mitochondrial reprogramming | emphysema | 2 | EMCN and epicardial progenitors in HUVECs |
| Kupffer cell | A Novel Missense Variant of ZC3H12A in Pulmonary Arterial Hypertension | PAH | 2 | CD20 and CD68 as B cell and macrophage markers |
| Mature luminal cell | CSF1R modulates megakaryopoiesis by targeting RUNX1 in immune thrombocytopenia | auto.-thromb. | 3 | CSF1R in megakaryocytes, platelets, and bone marrow |
| " | Molecular investigation and long-term clinical progress in Greek Cypriot families with recessive distal renal tubular acidosis and sensorineural deafness | RTA | 2 | ATP6V1B1 in intercalated and collecting duct cells |
| Hypercholesterolemia | Inflammation and Notch signaling: a crosstalk with opposite effects on tumorigenesis | JAG2 | 2 | LDLR–atherosclerosis link in cardiovascular context |
| Pulmonary fibrosis | Endothelial-derived PDGF-BB and HB-EGF coordinately regulate pericyte recruitment during vasculogenic tube assembly and stabilization | PDAP1 | 3 | PDGFB/PDGFRB/EGFR in vascular integrity and hemorrhage |
| " | The role of ADAM17 during liver damage | AREGB | 2 | AREG and MERTK roles in fibrosis |
| Renal tubular acidosis | Receptor-Mediated Endocytosis in the Proximal Tubule | OBP2A | 3 | Proximal tubule genes (OCRL, CTNS, CLCN5) that cause RTA |
| Rheumatoid arthritis | The microfibril-associated glycoproteins (MAGPs) and the microfibrillar niche | MFAP2 | 3 | TNF/TNFSF11/IL10 in inflammation and bone resorption |

Continued on next page

Continued from previous page

| Target topic | Source paper | From | Pairs | Cross-domain link |
|---|---|---|---|---|
| " | Macrophage-specific metalloelastase (MMP-12) truncates and inactivates ELR+ CXC chemokines and generates CCL2, -7, -8, and -13 antagonists | CXCL6 | 2 | MMP12 regulation of inflammation and leukocyte recruitment |
| ASIP | A distinct plasmablast and naïve B-cell phenotype in primary immune thrombocytopenia | activated B | 3 | BAFF binding to TACI, BAFF-R, and BCMA |
| CGA | Functional vulnerability of liver macrophages to capsules defines virulence of blood-borne bacteria | Kupffer cell | 4 | Receptor–ligand binding (ASGR, CRIg) in macrophage recognition |
| JAG2 | Intestinal Stem Cell Niche: The Extracellular Matrix and Cellular Components | intest. SC | 2 | Delta-like ligands activating Notch receptors in stem cell niche |
| PSEN1 | Hepatoblastoma: From Molecular Mechanisms to Therapeutic Strategies | hepatoblast | 18 | Notch pathway background lists all DLL/JAG–NOTCH LR pairs |

### S11 Configuration and reproducibility

#### S11.1 Configuration schema

Interaction-finder uses a hierarchical TOML configuration system. The configuration file specifies LLM models, search backends, and stage-specific parameters. The hierarchy for LLM model selection follows the fallback chain:

```
agents.module.agent -> agents.module._ -> agents._ -> code default
```

For example, `agents.extraction.judge` selects the model for the cross-document judgment agent, `agents.extraction._` sets the default for all extraction agents, and `agents._` sets the global default. This enables cost-quality tradeoffs: a cheap model for high-volume entity extraction, a more capable model for final judgment.

Example configuration:

```
[agents._]
llm = "openai:gpt-5-mini/low+flex"

[agents.extraction.judge]
llm = "openai:gpt-5/medium+flex"

[stage.keywords]
search_backend = "searxng"
document_context_chars = 50000

[stage.search]
search_backend = "pubmed"
```

#### S11.2 Checkpoint architecture

Self-contained JSON checkpoint files are produced at each pipeline stage, capturing the complete pipeline state: configuration, search results, extracted entities, pair assessments, and accumulated metadata. Checkpoints are a superset: each stage adds data while preserving all prior stage data.

This architecture provides:

- **Reproducibility**: checkpoints preserve all inputs, parameters, and intermediate results. Post-search stages operate on a fixed document set, reducing variability to LLM inference non-determinism.
- **Inspectability**: checkpoints are standard JSON, queryable with jq, loadable in pandas, or examinable in any JSON viewer.

- **Resumption**: if a run fails, it resumes from the last successful checkpoint.
- **Modularity**: stages can run independently. A researcher can run keyword extraction and search, inspect the discovered literature, and then run extraction on the same checkpoint.

#### S11.3 Managing non-determinism

LLM outputs are inherently non-deterministic: two runs from the same topic string may produce different results. Interaction-finder manages this in two ways:

1. **Search results are cached in checkpoints.** Once the search stages complete, the discovered literature is fixed. All subsequent extraction stages operate on this fixed document set, reducing variability to the LLM extraction and judgment steps.
2. **Baseline variability is quantified.** Each baseline was run 5 times per topic, producing error bars that characterise the expected variability of LLM-based extraction.

#### S12 Constructed negatives for plausibility scoring

The mixture model needs a reference distribution of implausible associations. We constructed one per domain using entities that are real biological entities (so the judge recognises them and engages with the pairing) but mechanistically unlikely to be associated with the topic. The three domains draw negatives from different sources, described below.

##### S12.1 Disease–gene

For each of the 20 disease topics, we identified the HPO branch (direct child of HP:0000118 "Phenotypic abnormality") containing the topic's HPO term, then selected the branch with the highest Jaccard dissimilarity in gene content. We collected all genes annotated to HPO terms in the distant branch, excluded any gene present in the topic's gold-standard set, and sampled at least 20 and up to the gold-standard count, capped by the number of genes available in the distant branch. For topics with very large gold-standard sets (e.g. dilated cardiomyopathy, 185 genes), the distant branch exhausts before matching the gold-standard count, yielding fewer negatives than positives; across the 20 topics this gives 918 scored negatives in total. HPO term-to-gene mappings were taken from the HPO `phenotype_to_genes.txt` release (v2025-05-06). Topic-to-HPO mappings were read from the case study term files used for evaluation.

##### S12.2 Celltype–cellmarker

The same branch-distance approach was applied using the Cell Ontology (CL, v2025-04-10), rooted at CL:0000000 ("cell"). CellMarker cell names were mapped to CL terms by label matching. For each topic, markers from the most dissimilar CL branch were collected and sampled as above.

##### S12.3 Ligand–receptor

Transcription factors are mechanistically incompatible with ligand–receptor binding at the cell membrane, making them strong negatives. We extracted human transcription factors (taxon 9606) from TFCheckpoint v2 (Acencio et al. 2024), excluded any that overlap with known receptors in the Ramilowski dataset (Ramilowski et al. 2015) (e.g. nuclear hormone receptors), and sampled 20 negatives per topic from the filtered set (gold-standard sets here contain only 1–8 receptors, so the minimum is binding).

### S13 Plausibility scoring protocol

Each candidate pair, plus a matched set of constructed negatives and the gold-standard reference for each topic, was scored for plausibility by an independent LLM judge with no access to the pipeline's evidence or reasoning. We used three scoring methods to check that the result is not an artefact of how plausibility is elicited, and report all three across the cross-method comparison below. The main manuscript reports Method B (direct 1–9 rating); Methods A and C are presented here as consistency checks. All methods used `temperature=0`. Pair counts depend on the positive reference set, discussed in Section S13.4 below.

#### S13.1 Scoring methods

The three methods probe plausibility at different levels: Method A decomposes the assessment into explicit dimensions, Method B captures a holistic judgement on a fine-grained scale, and Method C extracts the model's full probability distribution over ratings rather than a single greedy token.

**Method A: binary checklist (GPT-5).** The LLM answered six yes/no questions per pair (see system prompt below). Questions 1–5 are positive-direction; question 6 is negative-direction. The score is the proportion of answers indicating plausibility (yes for 1–5, no for 6), yielding 7 possible values on $[0, 1]$. This decomposition makes the basis for the score transparent but limits resolution to 7 levels.

**Method B: direct rating (GPT-5).** The LLM provided a single integer rating from 1 to 9 (see system prompt below), normalised to $[0, 1]$ as $(r - 1)/8$. The 9-point scale provides finer resolution than Method A and is the simplest method to explain.

**Method C: token probability (GPT-4.1).** The same prompt as Method B, but with `max_tokens=1` and `logprobs=true`. The score is the probability-weighted average over score tokens 1–9, normalised as in Method B. GPT-4.1 was used because GPT-5 does not expose token-level log-probabilities. The score incorporates probability mass across all nine rating levels rather than a single greedy choice, making it effectively continuous, but the change of model confounds the comparison with Methods A and B.

#### S13.2 Prompts

The user prompt was identical across methods:

```
Topic: {topic}
{type1}: {entity1}
{type2}: {entity2}
```

The order of the two entities was randomised per call to prevent position bias.

**System prompt (Method A):**

```
You are evaluating whether biological entity pairs represent plausible
associations. For each pair, answer these six questions with "yes" or "no"
and a one-sentence reason.

1. Are the two entities known to participate in the same biological
   pathway, process, or tissue context for the given topic?
2. Is there a known or plausible molecular mechanism by which one entity
   could directly affect or interact with the other?
3. Have the two entities been discussed together in peer-reviewed
   biomedical publications?
4. Is the proposed association consistent with the known molecular or
   cellular functions of both entities?
5. Would a biomedical researcher studying this topic consider the
   association worth investigating?
6. Is there a clear biological reason why these entities would NOT be
   associated in this context?

Respond in this exact format for each question:
1. yes/no — reason
2. yes/no — reason
...

Calibration examples:

  gene: BRCA1
  phenotype: breast cancer
  1. yes — (calibration)
  2. yes — (calibration)
  ...
  6. no — (calibration)

  gene: GJB2
  phenotype: emphysema
  1. no — (calibration)
  ...
  6. yes — (calibration)
```

**System prompt (Methods B and C):**

```
Rate the plausibility of biological entity-pair associations on a scale
of 1 to 9.
- 1: No plausible biological connection. A domain expert would not
  consider this worth investigating.
- 9: Well-established association with strong mechanistic basis.

Respond with the integer rating on its own line, then a one-sentence
reason.

Calibration examples:

  gene: BRCA1
  phenotype: breast cancer
  Rating: 9

  gene: GJB2
```

```
phenotype: emphysema
Rating: 1
```

The calibration examples shown above are for the disease–gene domain. Calibration pairs were chosen to be unambiguous: positives are well-established associations (BRCA1/breast cancer, CD3/T cell, VEGFA/KDR), and negatives pair real entities with no known biological connection in the relevant context. The celltype–cellmarker and ligand–receptor domains used domain-appropriate pairs:

**Table S19:** Calibration examples used per domain. One positive and one negative example were included in each prompt, in randomised order.

| Domain | Category | Entity 1 | Entity 2 |
|---|---|---|---|
| Disease–gene | Positive | BRCA1 | breast cancer |
| Disease–gene | Negative | GJB2 | emphysema |
| Celltype–cellmarker | Positive | CD3 | T cell |
| Celltype–cellmarker | Negative | Osteocalcin | T cell |
| Ligand–receptor | Positive | VEGFA | KDR |
| Ligand–receptor | Negative | VEGFA | TP53 |

#### S13.3 Per-topic score distributions

The aggregate distributions in Fig. 3 pool scores across 20 topics per domain. Fig. S7, Fig. S8, and Fig. S9 show the per-topic Method B distributions (with the all-gold positive reference used in the main manuscript; see Section S13.4) so the aggregate result can be checked against topic-level variation. The candidate–positive similarity holds across the majority of topics, but the candidate–negative separation is uneven: in disease–gene, a substantial minority of topics show candidate and negative distributions overlapping at the IQR level, with the overlap concentrated in topics whose curated gene lists are dominated by indirectly-supported associations (e.g. autoimmune thrombocytopenia, parkinsonism, pulmonary fibrosis, rheumatoid arthritis). In these topics the positive distribution is itself broad and reaches down toward the negative range, so the candidate distribution sits in the same region as both, and the aggregate gap visible in Fig. 3 is driven by the topics with cleaner literature support. The ligand–receptor reference sets are smaller still (half the topics have fewer than three gold-standard pairs); panels with fewer than three positives omit the gold-standard violin since a density estimate over so few points is not meaningful, and the underlying scores remain visible in the scatter column to the right of the violins.

#### S13.4 Positive reference set

The mixture model takes a positive distribution as a reference for what *real biological associations look like under the judge*. Two reasonable references are available, and each captures something different about the question being asked.

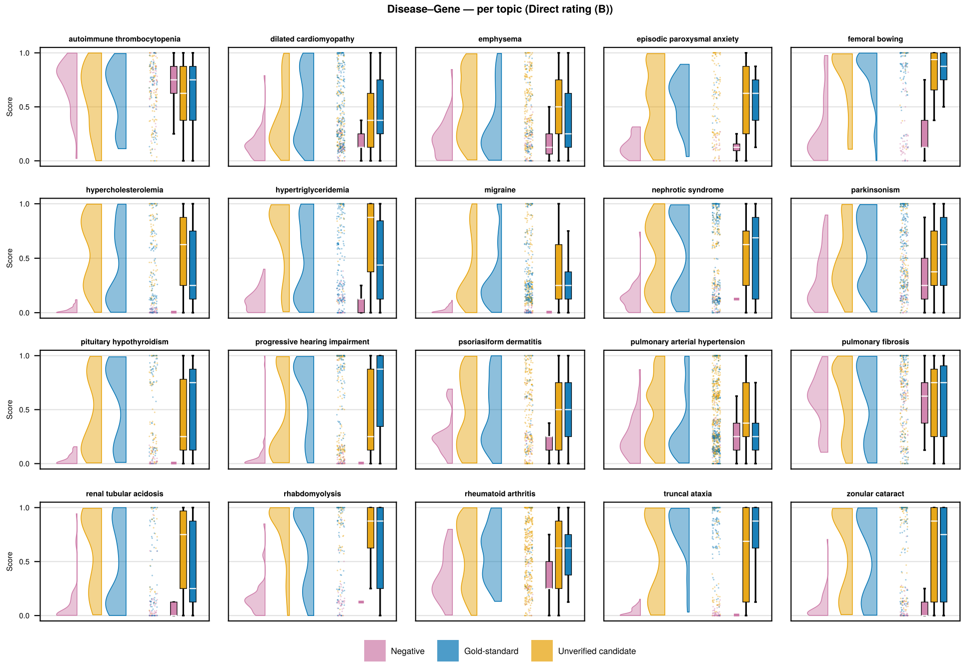

**Figure S7:** Per-topic plausibility score distributions for disease–gene (Method B). Each panel shows one of the 20 disease topics. Scores are shown for constructed negatives (purple), ambiguous candidates (orange), and gold-standard positives (blue). The candidate–positive similarity observed in the aggregate (Fig. 3) is consistent across the majority of individual topics.

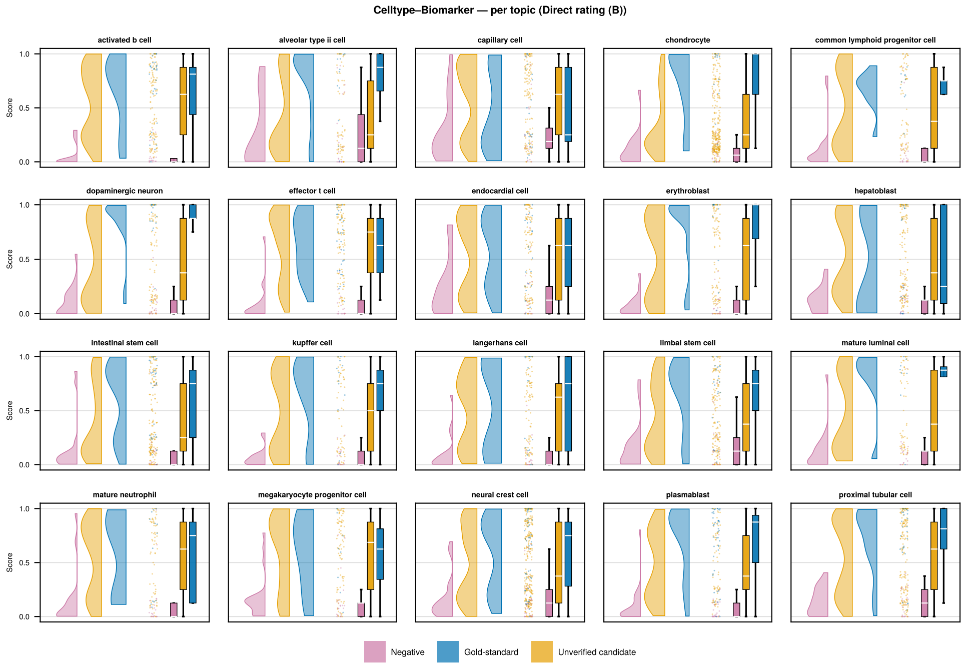

**Figure S8:** Per-topic plausibility score distributions for celltype–cellmarker (Method B). Each panel shows one of the 20 celltype topics. As in the disease–gene domain, candidate score distributions (orange) consistently resemble gold-standard positives (blue) rather than constructed negatives (purple), though topics with small gold-standard sets show wider positive distributions.

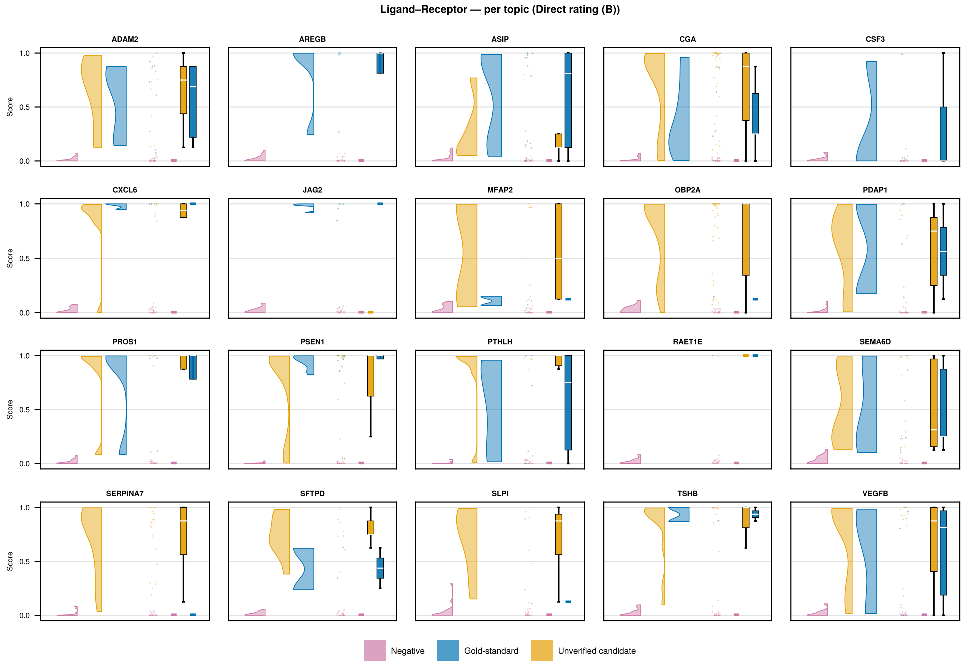

**Figure S9:** Per-topic plausibility score distributions for ligand–receptor (Method B). Each panel shows one of the 20 ligand topics. The reference sets are small (1–12 known receptors per ligand), and panels with fewer than three gold-standard positives omit the gold-standard violin; the individual scores remain visible in the scatter column.

The *retrieved* reference contains the gold-standard pairs that interaction-finder extracted from the literature and matched against the curated dataset. These are pairs for which direct literary support exists, the pipeline was able to find that support, and the LLM judge has training context for the relationship. The *all-gold* reference contains every pair in the curated dataset for each topic, regardless of whether interaction-finder recovered it. It includes pairs whose only documentation is indirect: for HPO, a (gene, phenotype) link inferred from a primary (gene, disease) annotation where the disease lists the phenotype as a clinical feature.

The retrieved reference is internally consistent, since every pair is one the judge can confidently score from training data, but it conditions on pipeline behaviour and so leaves the comparison vulnerable to the critique that candidates and gold pairs look alike only because both went through the same literary filter. The all-gold reference does not condition on pipeline behaviour, but the LLM judge under-rates the indirectly-supported pairs it adds, dragging the positive distribution into a range where it overlaps the constructed-negative distribution. Each reference answers a slightly different question, and neither is strictly better.

We report Method B against the all-gold reference as the headline in the main manuscript, on the grounds that not conditioning on pipeline behaviour is the more conservative choice when defending the claim that unverified candidates resemble real associations. The retrieved reference produces a more discriminative judge (higher AUC against constructed negatives) and is the like-for-like reference for Methods A and C below. Pair counts under each reference are in Table S20, and both references are reported in Table S21.

**Table S20:** Scored pair counts across 60 topics. The candidate and constructed-negative sets are identical across reference choices; only the positive set differs. The retrieved set counts gold-matching candidate rows the on-topic restriction retains: it removes a larger share of these rows (~47% for disease–gene) than of distinct gold partners (~9% overall, 590 of 649), because a dropped off-topic row's gold partner is often still recovered by another retained row.

| Domain | Positive (retrieved) | Positive (all-gold) | Constructed negative | Candidate |
|---|---|---|---|---|
| Disease–gene | 240 | 1114 | 918 | 1708 |
| Celltype–cellmarker | 122 | 411 | 414 | 1544 |
| Ligand–receptor | 10 | 80 | 400 | 58 |

#### S13.5 Cross-method comparison

With the reference choice settled, the question for the methods comparison is narrower: do Methods A and C reach the same qualitative conclusion as Method B? Table S21 reports AUC and $\hat{\pi}$ for all three methods on the retrieved reference (the like-for-like comparison), plus Method B at the all-gold reference for the headline.

All four configurations reliably separate gold-standard positives from constructed negatives (AUC 0.84–1.00), and $\hat{\pi}$ exceeds 0.6 in every condition with all 95% CIs excluding zero. The qualitative pattern is robust to both reference choice and scoring method.

**Table S21:** Discrimination (AUC) and estimated true-positive fraction ($\hat{\pi}$) across scoring methods and domains. All four configurations reliably separate gold-standard positives from constructed negatives. Method B (all gold) is the headline reported in the main manuscript; the retrieved rows enable a like-for-like comparison across the three methods. Method C uses GPT-4.1 due to log-probability availability; Methods A and B use GPT-5. 95% bootstrap CIs in brackets; /n/+ is the positive-set size used.

| Method | Domain | AUC | $\hat{\pi}$ [95% CI] | /n/+ |
|---|---|---|---|---|
| Direct rating, all gold (B) | Disease–gene | 0.835 | 0.79 [0.72, 0.83] | 1114 |
| | Celltype–cellmarker | 0.886 | 0.78 [0.70, 0.85] | 411 |
| | Ligand–receptor | 0.949 | 0.98 [0.92, 1.00] | 80 |
| Direct rating, retrieved (B) | Disease–gene | 0.890 | 0.63 [0.54, 0.69] | 240 |
| | Celltype–cellmarker | 0.908 | 0.72 [0.58, 0.81] | 122 |
| | Ligand–receptor | 0.999 | 0.91 [0.91, 0.91] | 10 |
| Checklist, retrieved (A) | Disease–gene | 0.855 | 0.71 [0.65, 0.77] | 240 |
| | Celltype–cellmarker | 0.895 | 0.75 [0.68, 0.79] | 122 |
| | Ligand–receptor | 0.946 | 0.97 [0.89, 1.00] | 10 |
| Token probability, retrieved (C) | Disease–gene | 0.917 | 0.97 [0.96, 0.99] | 240 |
| | Celltype–cellmarker | 0.895 | 0.99 [0.97, 1.00] | 122 |
| | Ligand–receptor | 0.951 | 1.00 [1.00, 1.00] | 10 |

On the retrieved reference, Method B produces the lowest $\hat{\pi}$ of the three methods, likely because its 9-point scale leaves room for intermediate scores that the mixture model assigns partially to the negative component. Method A, with only 7 score levels and a checklist structure that rewards partial matches, yields moderately higher estimates. Method C produces the highest and tightest estimates; the GPT-4.1 scores are notably less variable than GPT-5 scores, compressing most of the discrimination into fewer effective score levels and yielding narrower bootstrap intervals. How much of this is the token-probability method versus the model change is not separable here. Across all three methods, the score distributions (Fig. S10) show the same core pattern: candidates resemble gold-standard positives rather than constructed negatives.

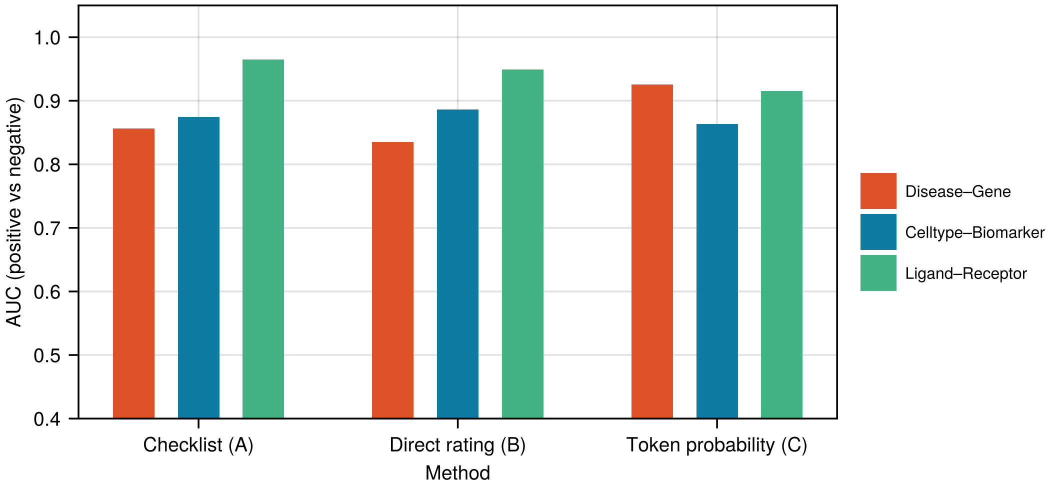

**Figure S10:** Plausibility score distributions for all three scoring methods (rows) across the three domains (columns). The pattern of candidates resembling gold-standard positives is consistent across methods.

#### S14 LLM prompts and output schemas

Each LLM agent in the pipeline receives a **system prompt** (defining its role and instructions) and a **user prompt** (providing the specific inputs for a given call). Outputs are constrained to structured JSON schemas via Pydantic models. All prompts and schemas are defined as module-level constants in the Python source; the canonical versions can be inspected in the repository. This section documents the prompts used in the evaluation. Seventeen agents across three stages cover the full pipeline (Table S22); two shared output schemas referenced by multiple agents are documented at the end.

**Table S22:** Roster of LLM agents and shared output schemas in the interaction-finder pipeline.

| Stage | Agent | Role |
|---|---|---|
| Keywords | Query expander | Generates queries targeting review articles for bridging-term mining. |
| | Document summariser | Summarises a review and extracts candidate bridging terms. |
| | Result selector (keywords) | Picks which review articles to fetch from a results page. |
| | Reflector (keywords) | Decides whether keyword coverage is sufficient to stop. |
| Search | Goal planner | Names subject areas the wide search should cover. |
| | Query generator | Writes broad/medium/focused/indirect queries against unmet goals. |
| | Result selector (wide search) | Selects relevant papers from each search round. |
| | Reflector (wide search) | Tracks goal coverage and decides whether to continue searching. |
| Extract | Document analysis | Scores paper quality and extracts entities of target types. |
| | Entity consolidation (pairwise) | Decides whether two similar entity names should merge. |
| | Entity consolidation (group/cluster) | Resolves clusters of similar names, including neighbour clusters. |
| | Relationship consolidation | Canonicalises relationship labels and classifies polarity. |
| | Proximal pair extraction | Extracts candidate associations from co-mention regions. |
| | Pair evidence judge | Scores per-document evidence quality for a candidate pair. |
| | Cross-document judge | Synthesises per-document evidence into accept/reject. |
| | Region assessment (co-mention sweep) | Assesses candidate pairs in regions surfaced by the co-mention sweep. |
| | Subject-trust judge | Marks whether a candidate's subject-side entity denotes the topic subject. |
| (schema) | `EvidenceQuality` | Shared evidence-quality schema (four factors + 1–9 score). |
| | `PaperQualityAssessment` | Shared seven-dimension paper quality schema. |

#### S14.1 Keyword discovery stage

The keyword discovery stage uses four LLM agents in an iterative loop: a query expander proposes review-article searches, a result selector picks results to fetch, a document summariser extracts bridging terms from each fetched review, and a reflector decides whether to continue or stop.

##### S14.1.1 Query expander

Generates search queries targeting review articles about the research topic.

**System prompt:**

```
You are an expert at generating search queries for finding
review articles and comprehensive summaries.

Your goal is to create queries that will find review articles,
meta-analyses, and comprehensive summaries about the given
topic. These articles should discuss the topic broadly and
mention related concepts that could serve as "bridging terms"
for literature search.

Guidelines:
- Target review articles, not primary research papers
- Include variations in terminology and synonyms
- Consider queries that would find papers discussing related
  areas and connections
- Keep queries concise but specific enough to find quality
  reviews
- Generate 1-5 queries, prioritizing quality over quantity

Focus on finding articles that will help identify bridging
terms: related concepts, alternative approaches, and connected
research areas that don't appear in the original topic name.

Explain your query strategy and why your queries will find
useful review articles.

Query format: {backend-specific directive}
```

**User prompt template:**

```
Generate search queries to find review articles about: {topic}
```

**Output schema** (`QueryExpansionOut`):

```
{
  "queries": ["<str>", ...],
  "reasoning": "<str>"
}
```

##### S14.1.2 Document summariser

Summarises each retrieved review article and identifies bridging terms: search keywords representing research contexts (disease subtypes, mechanisms, pathways) rather than specific entity instances.

**System prompt:**

```
You are an expert at summarizing scientific documents and
identifying bridging terms for literature search expansion.

Bridging terms are search keywords that help locate papers
from different angles, enabling search diversification. They
should represent research CONTEXTS (disease subtypes,
mechanisms, broad pathways, clinical presentations) rather
than specific entity instances being sought (individual genes,
proteins, markers, drugs, etc.).

Context terms help find varied papers, covering different
angles of the topic. Entity-specific terms just repeatedly
find papers about those same entities.

Given a document, provide:
1. A concise summary (50-750 chars)
2. Related research areas that directly inform the topic
3. Bridging terms (5-10 high-quality context terms)
```

**User prompt template:**

```
Review this document and identify HIGH-QUALITY bridging terms.

**Target topic:** {topic}

**Document content:**
{document_text}

**Extracted keywords (ranked by relevance to topic):**
{keywords_from_YAKE_RAKE_TFIDF_KeyBERT}
```

The user prompt includes detailed guidance on what constitutes a good bridging term, with positive and negative examples tailored to the topic's entity types.

**Output schema** (`DocumentSummaryOut`):

```
{
  "summary": "<str, 50-750 chars>",
  "related_areas": ["<str>", ...],
  "bridging_terms": ["<str>", ...],
  "coverage_contribution": "<str>"
}
```

##### S14.1.3 Result selector (keywords)

Selects which search results are most likely to be valuable review articles.

**System prompt:**

```
You are an expert at identifying review articles and
comprehensive summaries from search results.

Your task is to select which search results are most likely to
be valuable review articles that will help identify bridging
terms. Review the titles and snippets to make your selection.

Selection criteria:
- Prioritize review articles, meta-analyses, and systematic
  reviews
- Look for comprehensive discussions of the topic
- Prefer articles that discuss multiple aspects or
  perspectives
- Avoid primary research papers focused on narrow experimental
  results
- Look for articles that discuss related areas and connections

Return the indices of results to fetch, ordered by priority
(most valuable first). Typical selections are 5-10 results,
but adjust based on quality.
```

**User prompt template:**

```
Review these search results and select which ones to fetch
for keyword extraction.

Topic: {topic}

Search Results:
[0] {title}
{snippet}

[1] {title}
{snippet}
...

Select the indices of results that are most likely to be
valuable review articles.
```

**Output schema** (`ResultSelectionOut`):

```
{
  "selected_indices": [<int>, ...],
  "reasoning": "<str>"
}
```

##### S14.1.4 Reflector (keywords)

Assesses whether accumulated keyword coverage is sufficient or whether additional review articles should be sought.

**System prompt:**

```
You are an expert at assessing literature search coverage and
deciding when sufficient coverage has been achieved.

Given summaries of all documents processed so far, decide
whether to continue searching or stop.

Decision criteria for CONTINUE:
- Significant gaps in coverage remain
- Documents suggest important related areas not yet explored
- New perspectives or methodologies mentioned but not
  investigated
- Limited diversity in the documents found so far

Decision criteria for STOP:
- Comprehensive coverage of major aspects of the topic
- Diminishing returns (recent documents not adding much new)
- Good diversity of perspectives and approaches covered
- Sufficient bridging terms identified

If continuing, suggest new search angles based on gaps
identified in the coverage so far.

Be thoughtful but not overly perfectionistic. The goal is
reasonable coverage, not exhaustive coverage.
```

**User prompt template:**

```
Review the documents processed so far and decide whether to
continue searching.

Topic: {topic}
Current round: {n}/{max_rounds}
Documents processed: {count}

Document Summaries:
Document 1:
Summary: ...
Related areas: ...
Bridging terms: ...
Coverage: ...

Decide whether coverage is sufficient (stop) or more searches
are needed (continue).
```

**Output schema** (`ReflectionOut`):

```
{
  "decision": "continue" | "stop",
  "reasoning": "<str>",
  "new_search_angles": ["<str>", ...]
}
```

#### S14.2 Wide search stage

The wide search stage uses four LLM agents in a search–reflect loop: a goal planner names the subject areas the search should cover, a query generator writes queries at four complexity levels against unmet goals, a result selector picks the relevant papers from each round, and a reflector updates goal coverage and decides whether to stop.

##### S14.2.1 Goal planner

Identifies subject areas and research domains that should be covered during the search.

**System prompt:**

```
You are an expert research strategist planning comprehensive
literature searches.

Your task is to identify subject areas and research domains
that should be covered to ensure comprehensive literature
discovery on a given topic.

Guidelines for Comprehensive Coverage:
- Consider the core topic and its direct applications (basic
  and clinical research)
- Identify related methodologies, techniques, and approaches
  (experimental, computational, clinical)
- Think about adjacent research areas and intersecting domains
- Consider both theoretical foundations (mechanisms, pathways)
  and practical applications (diagnostics, therapeutics)
- Include areas that might use different terminology for
  similar concepts
- Consider different research perspectives: molecular,
  cellular, systems-level, clinical, translational
- Think broadly about what researchers in this field study:
  genetics, epigenetics, signaling, metabolism, imaging,
  biomarkers
- Include both well-established areas and emerging/novel
  research directions
- Aim for 8-15 well-defined subject goals that span the
  research landscape

Be ambitious about coverage - it's better to identify more
goals that guide thorough exploration than to miss important
research areas. Your goals should guide query generation to
```

```
ensure diverse, comprehensive coverage without drifting into
irrelevance.
```

**User prompt template:**

```
Research topic: {topic}

Keyphrases available: {keyphrase1}, {keyphrase2}, ...

Identify subject areas and research domains that should be
covered to ensure comprehensive literature discovery.
```

**Output schema** (`SubjectGoalsOut`):

```
{
  "goals": ["<str>", ...],
  "reasoning": "<str>"
}
```

##### S14.2.2 Query generator

Generates search queries at four complexity levels targeting unsatisfied subject goals.

**System prompt:**

```
You are an expert at generating search queries for academic
literature discovery.

Your task is to create diverse, targeted search queries that
will find relevant papers and articles. You will be given:
- A research topic
- A list of keyphrases to incorporate
- Subject goals to target (unsatisfied areas needing coverage)

REQUIRED: Generate queries at ALL FOUR complexity levels
below. Each level serves a distinct purpose:

1. BROAD queries (1-2 general concepts, 3-6 words) -- minimum
   2 required:
   - Combine the main topic with ONE general domain concept
   - NO specific named entities beyond the topic itself
   - Purpose: Cast a wide net, ensure baseline coverage

2. MEDIUM queries (2-3 concepts, 6-10 words) -- minimum 2
   required:
   - Combine specific mechanisms with conditions using natural
     phrases
   - Use at most ONE specific entity per query
   - Purpose: Target specific but well-documented areas

3. FOCUSED queries (3+ concepts, 10-15 words) -- minimum 2
   required:
   - Highly specific research questions with clear semantic
```

```
    structure
  - Limit to 1-2 specific entities; prefer process/mechanism
    names over entity lists
  - Purpose: Find specialized literature

4. INDIRECT queries (without main topic subject) -- minimum 2
   required:
   - Completely OMIT the main research topic; use only related
     keyphrases, mechanisms, or phenotypes
   - Include both broad and focused indirect queries
   - Purpose: Find papers discussing relevant biology without
     explicit topic mention

Query Generation Guidelines:
- Structure queries as natural phrases with connecting words,
  not keyword concatenation
- Prioritize breadth first: cast a wide net with broad queries
  before diving deep
- Target unsatisfied subject goals explicitly
- Incorporate provided keyphrases naturally but don't force
  all keyphrases into every query
- Use variations in terminology, synonyms, and related
  concepts
- Avoid redundancy with previous queries

Query format: {backend-specific directive}
```

**User prompt template:**

```
Research topic: {topic}

Keyphrases to incorporate: {keyphrase1}, {keyphrase2}, ...

Subject goals (unsatisfied): {goal1}, {goal2}, ...

Round {n} of {max_rounds}

Generate search queries that target unsatisfied subject goals
and incorporate the keyphrases.
```

**Output schema** (`QueryGenerationOut`):

```
{
  "broad_queries": ["<str>", ...],
  "medium_queries": ["<str>", ...],
  "focused_queries": ["<str>", ...],
  "indirect_queries": ["<str>", ...],
  "reasoning": "<str>"
}
```

##### S14.2.3 Result selector (wide search)

Selects relevant papers from each search round with an inclusive strategy.

**System prompt:**

```
You are an expert at identifying relevant research papers for
comprehensive literature collection.

Your task is to select search results that are relevant to the
research topic. The goal is to build a broad collection of
papers for downstream entity extraction and analysis, so adopt
an inclusive selection strategy.

Result quality varies widely by batch: some searches return
highly relevant results requiring selection of most or all
papers; other searches yield few or no relevant papers
requiring minimal selection. Judge each batch independently
based on actual relevance rather than targeting a fixed
percentage.

Selection Strategy -- Include results that meet ANY of these
criteria:
- Directly relevant to the research topic
- Discusses mechanisms, pathways, biological processes, or
  clinical applications related to the topic
- Reports primary research findings ON the topic
- Provides overviews of the topic or related domains
- Examines methodologies, techniques, or approaches used to
  study the topic
- Discusses entities or associations that plausibly connect to
  the topic

When uncertain about a result's relevance based solely on
title and snippet, include it rather than exclude it.
Downstream processing will filter for quality and extract
specific information.

What to Exclude -- Exclude only results that are:
- Clearly off-topic or unrelated to the research domain
- Duplicate content
- Non-academic content with no substantive research discussion

Coverage Summary -- After making your selection, provide a
summary that includes:
- What subject areas and research domains are covered by
  selected results
- What types of research are represented
- What specific aspects of the topic these results address
- Any notable entities, pathways, or connections mentioned

Your selection builds the literature collection for entity
extraction. Prioritize breadth and recall over precision.
```

**User prompt template:**

```
Research topic: {topic}

Subject goals: {goal1}, {goal2}, ...

Search results from round {n}:
```

```
[0] {title}
{snippet}
URL: {url}

[1] {title}
...

Select the most relevant results and summarize what subject
areas they cover.
```

**Output schema** (`ResultSelectionOut`):

```
{
  "selected_indices": [<int>, ...],
  "covered_topics_summary": "<str>",
  "reasoning": "<str>"
}
```

##### S14.2.4 Reflector (wide search)

Evaluates coverage against subject goals and decides whether to continue or stop searching.

**System prompt:**

```
You are an expert at evaluating literature search coverage and
deciding when sufficient breadth has been achieved.

Your task is to reflect on search results collected so far and
decide whether to continue searching or stop.

You will be given:
- The research topic
- Original subject goals to cover
- Currently satisfied goals
- Summaries of search results from each round

Evaluation criteria:
- Assess which subject goals are now well-covered (mark as
  satisfied)
- Identify any new important subject areas discovered that
  should be explored
- Determine if another search round would likely add
  significant value
- Consider the quality and diversity of results found so far
- Balance comprehensiveness with diminishing returns

Decision guidelines for WHEN TO CONTINUE:
- Continue if ANY major subject goals remain unsatisfied
- Continue if new important areas were discovered that warrant
  exploration
- Continue if fewer than 50-75 unique results have been
  collected
- Continue if recent rounds are still finding diverse,
```

```
  non-repetitive results
- Continue if the variety of result types is limited
- Favor thoroughness - it's better to do one extra round than
  to stop prematurely

Decision guidelines for WHEN TO STOP:
- Stop if nearly all subject goals are well-covered AND
  sufficient result volume has been achieved
- Stop if results are becoming highly repetitive across
  multiple rounds with little new content
- Stop if the last 2 rounds added minimal new diverse results
  despite different queries
- Stop if we've reached the maximum configured search rounds

IMPORTANT: Require strong evidence before stopping. "Adequate
coverage" is not sufficient - aim for "comprehensive coverage"
with diverse result types and sufficient volume (aim for 50+
unique results for most research topics). Be thorough rather
than conservative.
```

**User prompt template:**

```
Research topic: {topic}

Subject goals: {goal1}, {goal2}, ...

Satisfied goals: {satisfied_goal1}, ...

Search summaries so far:
Round 1: {coverage_summary}
Round 2: ...

Current round: {n} of {max_rounds}

Evaluate coverage and decide whether to continue or stop.
```

**Output schema** (`ReflectionOut`):

```
{
  "satisfied_goals": ["<str>", ...],
  "new_goals": ["<str>", ...],
  "should_continue": <bool>,
  "reasoning": "<str>"
}
```

#### S14.3 Extraction stage

The extraction stage uses nine LLM agents working roughly in pipeline order. The document analyser scores paper quality and extracts entities from each retrieved document. Two consolidation agents merge entity names across documents: a pairwise consolidator for binary decisions on similar pairs, and a cluster consolidator for multi-member clusters and the neighbour-context pass. A proximal pair extractor proposes candidate associations from co-mention regions, and a region assessor checks co-occurrences missed by the proximal pass. A pair evidence judge scores per-document evidence for each candidate, while a relationship consolidator canonicalises the relationship labels. A cross-document judge synthesises everything into a final accept/reject decision. Finally, a subject-trust judge marks each accepted pair as on- or off-subject, supplying the report's default filter.

##### S14.3.1 Document analysis

Combined paper quality assessment (seven-dimension rubric, Section S5) and entity extraction in a single pass. Quality assessment comes first to establish paper understanding before extraction.

**System prompt:**

```
You are an expert biomedical scientist skilled in both
methodological critique and entity recognition.

Your task has TWO parts, completed in order:

PART 1: Paper Quality Assessment

Evaluate the paper's quality using the seven-dimension rubric
in the output schema.

Calibration:
- Score 2 is typical for solid published papers with standard
  reporting
- Score 0 and 3 are reserved for clear cases (obvious
  problems or exemplary practice)
- When uncertain between adjacent scores, prefer the lower one
- Reference specific observations from the text in each
  justification

If key sections (e.g., Methods) are absent, note this and
score based on what IS present.

PART 2: Entity Extraction

Identify and extract biological entities from the scientific
text.

Extraction rules:
1. Extract only entities of the requested types (gene,
   disease, protein, etc.)
2. Use canonical names (e.g., "BRCA1" not "BRCA-1")
3. Include all verbatim names as they appear in the text
4. Provide exact quotes supporting each entity
5. Only extract entities clearly relevant to the topic
6. Require clear textual support for every entity
7. Explain your reasoning for each entity extraction

Alias handling:
- Aliases must be alternative names for the SAME entity
- Include acronyms, abbreviations, and lexical variants
```

```
- If a name includes the base entity PLUS additional
  qualifiers that significantly narrow or change the meaning,
  extract it as a separate entity instead

If no entities are found: Return an empty entities list.
```

**User prompt template:**

```
# Document
**Title:** {title}

{document_text}

# Task
1. First, assess paper quality using the seven-dimension rubric
2. Then, extract entities relevant to: {topic}

**Target entity types:** {type1}, {type2}

For each entity of the specified types relevant to the topic,
provide:
- Canonical name
- All verbatim name variants from text
- Supporting quotes
- Reasoning for inclusion
```

**Output schema** (`DocumentAnalysisOut`):

```
{
  "paper_quality": {
    "method_clarity":          {"score": <0-4>, "justification": "<str>"},
    "data_provenance":         {"score": <0-4>, "justification": "<str>"},
    "statistical_rigour":      {"score": <0-4>, "justification": "<str>"},
    "internal_consistency":    {"score": <0-4>, "justification": "<str>"},
    "plausibility":            {"score": <0-4>, "justification": "<str>"},
    "reproducibility_signals": {"score": <0-4>, "justification": "<str>"},
    "integrity_indicators":    {"score": <0-4>, "justification": "<str>"}
  },
  "entities": [{
    "kind": "<str>",
    "name": "<str>",
    "aliases": ["<str>", ...],
    "quotes": ["<str>", ...],
    "reasoning": "<str>"
  }, ...]
}
```

##### S14.3.2 Entity consolidation (pairwise)

Decides whether pairs of similar entity names should be merged, renamed, or kept separate.

**System prompt:**

```
You are an expert at resolving entity naming ambiguities for
biomedical literature mining.

Context: You are given pairs of entities where one name
contains or is similar to the other. Each pair shows:
- A child term (the longer/more specific name)
- A candidate parent it might merge into (the shorter/broader
  name)

You will be provided a research topic and target entity types.
Use the topic to judge whether distinctions matter.

Only return pairs that should merge or be renamed. Pairs you
omit will be kept separate.

Actions:

merge (set rename=null) -- Merge the child into the parent
when the child is the same entity with a redundant qualifier
or variant.
Examples:
  'p53 gene' -> 'p53' -- merge ("gene" is redundant)
  'mutant BRCA1' -> 'BRCA1' -- merge (variant of same gene)
  'human insulin' -> 'insulin' -- merge (species qualifier)

rename (set rename="standard name") -- Replace verbose
descriptions with standard names, or normalize overly-specific
variants to a topic term when the extra specificity doesn't
add meaningful distinction for this research.
Examples:
  'transforming growth factor beta protein' -> 'growth factor'
    -- rename="TGF-beta"
  'mitogen-activated protein kinase enzyme' -> 'kinase'
    -- rename="MAPK"
  'familial idiopathic pulmonary arterial hypertension'
    -> 'hypertension' -- rename="pulmonary arterial
    hypertension" (normalize to topic)
Only rename to widely recognized standard names or to a topic
term when appropriate.

omit (do NOT return the pair) -- Keep both entities separate
when they represent genuinely distinct concepts relevant to
the research topic.
Examples:
  'MAP kinase' -> 'kinase' -- omit (MAPK is a specific family)
  'adrenergic receptor' -> 'receptor' -- omit (specific type)
  'p53 pathway' -> 'p53' -- omit (gene vs pathway)
```

**User prompt template:**

```
**Research topic:** {topic}

**Target entity types:** {type1}, {type2}

**Entity pairs to evaluate:**
1. [xK7m] 'p53 gene' -> 'p53' ?
2. [aB3c] 'TGF-beta protein' -> 'growth factor' ?
```

```
...

Only return pairs that should merge or be renamed.
Omit pairs that should remain separate.
```

Each pair includes a random 4-character confirmation token to prevent entity name hallucination during structured output generation.

**Output schema** (`EntityConsolidationDecisions`):

```
{
  "decisions": [{
    "pair_id": <int>,
    "confirm_token": "<4 chars>",
    "rename": "<str>" | null,
    "reasoning": "<str>"
  }, ...]
}
```

##### S14.3.3 Entity consolidation (group/cluster)

Reviews clusters of similar entities for potential merging. Used in both the main entity consolidation pass and the neighbour-based consolidation pass.

**System prompt:**

```
You are an expert at consolidating biomedical entity names for
systematic literature analysis.

You will be given groups of entities that all connect to the
same anchor entity. For each group, decide if all members
should unify to a single canonical name, based on the research
topic.

Decision framework:
1. Evaluate each group in the context of the research topic
2. Ask: Are these the same entity, or subtypes that the
   research topic wouldn't distinguish between? If so, merge.
   If the distinction matters for this research, reject.
3. Apply these principles:
   - Different phrasings of the same concept -> merge
   - Variant spellings or abbreviations -> merge
   - Subtypes of a concept the topic targets -> merge to that
     concept
   - Entities whose distinction matters for the research
     -> reject
   - One member doesn't belong -> exclude it

Actions:
- merge: Members should unify -> specify target (member number,
  member name, or a new parent concept). When choosing a merge
  target, select the most specific term that accurately
```

```
  encompasses all members.
- reject: Members are distinct entities that coincidentally
  share words or an interaction partner -> keep them all
  separate
- exclude: One specific member doesn't belong -> specify which
  one to remove
- split: Cluster mixes unrelated entities -> system splits at
  weakest link

You must return an explicit decision for every group.
```

**User prompt template (main consolidation):**

```
**Research topic:** {topic}
**Target entity types:** {type1}, {type2}

**Entity groups to consolidate:**

## Group xK7m
Members:
  1. interleukin 6
  2. IL-6
  3. IL6
```

**User prompt template (neighbour consolidation):**

```
**Research topic:** {topic}

**Neighbour clusters to evaluate:**

## Group aB3c
**Context:** These entities all connect to **chondrocyte** (cell)

Members:
  1. collagen type II
  2. COL2A1
  3. type II collagen
```

**Output schema** (`ClusterDecisions`):

```
{
  "decisions": [{
    "group_id": "<str>",
    "action": "merge" | "reject" | "split" | "exclude",
    "target": "<str>" | null,
    "reasoning": "<str>"
  }, ...]
}
```

##### S14.3.4 Relationship consolidation

Consolidates synonymous relationship labels, classifies polarity, and detects opposing relationships.

**System prompt:**

```
You are an expert at consolidating and classifying biological
relationship labels for literature mining.

Your task: For each relationship label, determine:
1. CONSOLIDATION: Should it be mapped to a canonical form?
2. POLARITY: What is its biological direction/effect?
3. OPPOSITES: Which other relationships have opposite
   biological effects?

PART 1: Consolidation Guidelines

Merge synonymous labels:
- "linked_to", "connected_to", "related_to"
  -> "associated_with"
- "upregulates", "increases expression of" -> "activates"
- "downregulates", "decreases expression of" -> "inhibits"

Preserve distinct biological meanings:
- "activates" vs "inhibits" (opposite effects)
- "regulates" vs "activates" (general vs specific)
- "binds_to" vs "activates" (physical vs functional)
- "causes" vs "associated_with" (causal vs correlational)

Standardize to common forms:
- Prefer active voice: "activates" over "is activated by"
- Prefer standard terms: "associated_with" over "linked_to"
- Prefer verbs: "regulates" over "regulation_of"

Ensure labels are short verb phrases without specific
subjects: 1-2 word verb phrase, generally applicable.

PART 2: Polarity Classification

Classify each relationship by its biological polarity:

POSITIVE - Promoting/increasing/activating effect
  Examples: "causes", "activates", "promotes", "upregulates"

NEGATIVE - Inhibiting/decreasing/protective effect
  Examples: "protects_against", "inhibits", "prevents",
  "downregulates", "treats"

NEUTRAL - Mechanistic but directionality is ambiguous
  Examples: "regulates", "interacts_with", "binds_to",
  "associated_with"

IRRELEVANT - Orthogonal to biological mechanism
  Examples: "spatial_colocalization", "mentioned_together"

PART 3: Opposition Detection

Identify relationships with opposite biological effects:

Direct opposites:
- "activates" <->"inhibits"
```

```
- "increases_risk_of" <->"protects_against"
- "promotes" <->"prevents"
- "causes" <->"treats" (in disease context)

Non-opposites (do NOT mark):
- Different specificity levels: "regulates" is NOT opposite to
  "activates"
- Different mechanisms: "binds_to" is NOT opposite to
  "inhibits"
- Neutral vs directional: "associated_with" has no opposites

Important Guidelines:
- Consolidated labels MUST be 1-2 words maximum
- If label is already canonical, use it unchanged
- Multiple originals can map to same consolidated label
- Empty opposites list is valid and common
- When uncertain about directionality, prefer neutral over
  irrelevant
```

**User prompt template:**

```
**Research topic:** {topic}

**Target entity types:** {type1}, {type2}

**Relationship labels found (with frequency):**
- 'associated_with' (42)
- 'activates' (17)
- 'linked_to' (8)
...

For each relationship, provide:
1. Consolidated canonical form
2. Polarity classification relative to this research topic
```

**Output schema** (`RelationshipConsolidations`):

```
{
  "consolidations": [{
    "original": "<str>",
    "consolidated": "<str>",
    "polarity": "positive" | "negative" | "neutral" | "irrelevant",
    "opposites": ["<str>", ...],
    "reasoning": "<str>"
  }, ...]
}
```

##### S14.3.5 Proximal pair extraction

Identifies entity-entity associations within localised text regions where multiple entities co-occur.

**System prompt:**

```
You are an expert at identifying biological associations in
scientific text.

Your task: extract binary relationships between biological
entities from a localized text region.

You will be given:
1. A text region where multiple entities co-occur
2. A list of entities (with canonical names and aliases) to
   look for
3. The research topic for context

Extraction rules:
1. Only extract associations between entities in the provided
   list
2. Extract relationships that are clearly stated or strongly
   implied in the text
3. Use canonical entity names (not aliases) in your output
4. Each pair connects exactly two entities (binary
   relationships)
5. Provide exact quotes supporting each association
6. Suggest multiple relationship types if text implies
   different aspects (as separate list items)

Relationship type guidelines:
Each label must be SHORT (1-2 words), a verb phrase, generally
applicable across any entity pair.
- GOOD: ["regulates", "activates"], ["mutated_in"], ["treats"]
- BAD: ["promotes / enhances"], ["promotes (dysfunction)"]
- Multiple aspects? -> Multiple list items: ["promotes",
  "activates"] NOT ["promotes / activates"]

Standard relationship types (prefer these):
- Positive: "activates", "promotes", "upregulates", "causes"
- Negative: "inhibits", "prevents", "downregulates", "treats",
  "contraindicates"
- Neutral: "regulates", "interacts_with", "binds",
  "mutated_in", "associated_with", "no_effect"

Both positive and negative relationships are valuable -
inhibitory effects, contraindications, and explicit "no
effect" findings are just as important as activating
relationships.

Quality standards:
- Be conservative: only extract well-supported associations
- Quotes must directly support the claimed relationship
- Do not infer relationships from separate mentions without
  connecting evidence
- Co-occurrence alone is not sufficient - there must be
  stated/implied connection
- Prefer specific relationship types over generic
  "associated_with"

Important:
- Use canonical entity names exactly as shown (in bold)
- If no associations are found, return an empty pairs list
```

```
  with explanation in reasoning
- Always provide reasoning for your extraction choices
```

**User prompt template:**

```
# Context
Extract entity associations from this text region.

**Topic:** {topic}

**Entities in this region:**
- **{entity1_name}** [kind: {entity1_kind}, aliases: {entity1_aliases}]
- **{entity2_name}** [kind: {entity2_kind}, aliases: {entity2_aliases}]
- ...

# Document Extract
{text_region}

# Output
For each binary association between these entities that is
clearly stated or implied:
- Use the canonical entity names as shown in bold
- Provide exact supporting quotes
```

**Output schema** (`ProximalPairExtraction`):

```
{
  "pairs": [{
    "entity1": "<str>",
    "entity2": "<str>",
    "relationship_types": ["<str>", ...],
    "supporting_quotes": ["<str>", ...]
  }, ...],
  "reasoning": "<str>"
}
```

##### S14.3.6 Pair evidence judge

Assesses the strength of evidence for a specific entity pair within a single document.

**System prompt:**

```
You are an expert at evaluating evidence for biological
associations in scientific text.

Your task is to assess the strength of evidence for a specific
entity-entity association in a single document and select the
most appropriate relationship type.

You will be given:
1. A pair of entities (entity1 and entity2)
2. Candidate relationship types suggested from the text
3. Relevant text containing quotes supporting this association
```

```
4. The research topic for context

Evidence assessment:
Assess the four observable factors (directness, source_type,
specificity, language) based on what you observe in the text,
then assign an evidence_level consistent with those factors.

Relationship selection:
Select the SINGLE most accurate relationship type:
1. Prefer specific over generic (e.g., "activates" >
   "regulates" > "associated_with")
2. Consider directionality (e.g., "activates" vs "inhibits"
   vs "regulates")
3. If candidates are too generic, suggest a more specific type
   from the text
4. If multiple types are equally valid, choose the one with
   strongest evidence

Standard types: regulates, upregulates, downregulates,
activates, inhibits, interacts_with, binds, causes, prevents,
treats, mutated_in, associated_with

Critical:
- Assess factors first, then derive evidence_level from them
- Relationship type must be SHORT (1-2 words), a verb phrase,
  generally applicable
- Provide detailed reasoning referencing specific evidence
- Reference quote indices that most strongly support your
  assessment
```

**User prompt template:**

```
# Context
Assess evidence for an entity association.

**Topic:** {topic}

**Pair:** {entity1_name} ({entity1_kind}) <-> {entity2_name} ({entity2_kind})

**Relationship type candidates:** {candidates}

**Supporting quotes:**
{quotes}

# Document Extract
{text_region}

# Output
- Select the most appropriate relationship type
- Assess evidence quality factors and overall level
- Explain your reasoning
```

**Output schema** (`PairEvidenceJudgment`):

```
{
  "supporting_quote_ids": [<int>, ...],
  "reasoning": "<str>",
```

```
  "relationship": "<str>",
  "evidence": <EvidenceQuality>,
  "topic_relevance": <1-5>
}
```

##### S14.3.7 Cross-document judge

Makes a final accept/reject decision by synthesising evidence across all documents for a given entity pair. Two prompt variants handle unidirectional and contentious (contradictory evidence) pairs.

**System prompt:**

```
You are an expert scientific reviewer synthesizing evidence
across multiple documents.

Your task is to make a final judgment on whether an
entity-entity association is valid based on evidence from
multiple sources. You will receive:
1. Per-document assessments with evidence quality factors and
   reasoning
2. All supporting quotes from all documents
3. The topic and entity information

Decision framework:
- Accept when overall >= 6 across multiple sources OR single
  source with level >= 8
- Accept with caution when overall 5-6 with consistent
  supporting evidence
- Reject when overall <= 4 across sources OR contradictory
  high-level evidence

Evidence synthesis:
Synthesize the per-document evidence factors into an overall
assessment:
- directness: Use the most direct evidence available across
  documents
- source_type: Primary research takes precedence over reviews
- specificity: Prefer mechanistic over associative when
  available
- language: Note if sources use consistent or conflicting
  certainty
- overall: Weight by source quality and consistency

Key considerations:
1. Consistency: Do sources agree on the relationship nature?
2. Independence: Multiple independent sources vs. citations of
   same work?
3. Quality: Primary research > reviews > commentary
4. Mechanism: Is there explanation of how the relationship
   works?
5. Contradictions: How to weigh conflicting evidence?

Special cases:
```

```
- Conflicting biological effects (e.g., "activates" vs
  "inhibits") -> may indicate context-dependent effects;
  investigate carefully
- Single high-level source (8-9) + no other evidence -> accept
  cautiously (level ~7)
- All low-level (<=4) -> reject unless consistently suggestive

Decision guidance:
- Make a clear accept/reject decision
- Synthesize evidence factors across documents
- Relationship label must be SHORT (1-2 words), a verb phrase,
  generally applicable
- Explain what tipped the balance in your reasoning

Decision probability (decision_confidence):
After completing your analysis, estimate the probability that
your accept/reject decision is correct.

Calibration anchors:
- 0.95: Near-certain. Multiple independent high-quality
  sources agree; no reasonable doubt
- 0.85: Confident. Strong evidence with minor gaps
- 0.75: Probable. Good evidence, but some ambiguity
- 0.65: Lean. Evidence points one way but alternative
  interpretation exists
- 0.55: Slight lean. Marginal evidence; could go either way
- 0.50: Coin flip. Genuinely uncertain

When uncertain between two probability levels, prefer the
lower one.
```

**User prompt template (unidirectional):**

```
# Context
Make a final judgment on an entity association.

**Topic:** {topic}

**Pair:** {entity1_name} <-> {entity2_name}

**Relationship types found across documents:** {relationships}

# Document Extracts

## Document extract [1_abc12345]: {title}
**Assessment:** level 8/9 - causes
**Factors:** explicit, primary, mechanistic, definitive
**Reasoning:** ...
{text_region}

## Document extract [2_def67890]: {title}
...

# Task
Synthesize the evidence across documents.
```

For contentious pairs with contradictory evidence (e.g. both “activates” and “inhibits”), the prompt separates evidence by polarity (positive, negative, neutral) and asks the model to weigh the contradictions.

**Output schema** (`CrossDocumentJudgment`):

```
{
  "reasoning": "<str>",
  "relationship": "<str>",
  "evidence": <EvidenceQuality>,
  "accepted": <bool>,
  "decision_confidence": <0.0-1.0>,
  "topic_relevance": <1-5>
}
```

##### S14.3.8 Region assessment (co-mention sweep)

Assesses candidate entity pairs in text regions identified by the co-mention sweep pass (Section S1.3, step 6).

**System prompt:**

```
Analyze co-occurring entity mentions and assess their
relationships.

Task:
You will be given a text region and a list of candidate entity
pairs to evaluate. For each pair:
1. Verify that both entity mentions in the text actually refer
   to the specified canonical entities (not similarly-named
   entities or false matches)
2. If both entities are valid and relevant to the research
   topic, determine if the text makes any claim about their
   relationship

Only include a pair in pairs if BOTH conditions are met:
- Both entity mentions are valid (refer to the specified
  entities and are topic-relevant)
- The text states or strongly implies a relationship between
  them

Entity validation:
Check whether each mention refers to the specified canonical
entity:
- "BRCA1" vs "BRCA1-like protein" are different entities
- An entity mentioned in an unrelated context should not be
  included
- If either entity in a pair fails validation, do not include
  the pair

Relationship types:
Use one of the known relationship types listed in the prompt
when possible. If none fit, use a concise descriptive label.
```

```
Both positive and negative relationships matter - inhibitory
effects, contraindications, and explicit "no effect" findings
are just as important as activating relationships.

Quality standards:
- Be conservative: only report relationships that are clearly
  stated or strongly implied
- Co-occurrence alone is NOT sufficient - there must be a
  stated connection
- The relationship must be about these specific entities, not
  general statements
- Provide exact verbatim quotes from the text, not paraphrases
- If no pairs meet the criteria, return an empty pairs list
```

**User prompt template:**

```
# Context
Evaluate candidate entity pairs for associations in this
text region.

**Topic:** {topic}

**Candidate pairs to check:**
- {entity1_name} ({entity1_kind}) <-> {entity2_name} ({entity2_kind})
- ...

**Known relationship types:**
{known_relationships}

# Document Extract
{text_region}

# Task
For each candidate pair, determine if there is evidence
of a relationship in the text.
Provide supporting quotes for confirmed relationships.
```

**Output schema** (`RegionAssessmentOut`):

```
{
  "pairs": [{
    "entity1_name": "<str>",
    "entity2_name": "<str>",
    "relationship": "<str>",
    "reasoning": "<str>",
    "topic_relevance": <1-5>,
    "supporting_quotes": ["<str>", ...],
    "evidence": <EvidenceQuality>
  }, ...]
}
```

##### S14.3.9 Subject-trust judge

Marks each accepted pair as on- or off-topic (Section S1.4). One call per candidate subject-side entity, on GPT-5-mini. The subject anchor and kind are supplied directly, and the judge reasons only about the taxonomic relation between the candidate and the subject.

**System prompt:**

```
You are vetting a TRUSTED answer set for a research topic. You are given a
SUBJECT and a CANDIDATE, both of the same KIND. Decide whether the candidate
denotes the subject — the same entity, or a taxonomic form or category of it —
as opposed to a distinct entity that is merely related to it. Judge ONLY the
taxonomic relationship between the two; never reason about what either is
associated with, produces, expresses, binds, or is marked by.

ADMIT only when the candidate is one of:
  - the subject itself — an alias, acronym, synonym, or the same entity under a
    different name, stage, or form;
  - a more specific kind of the subject — a subtype, form, variant, or named
    member that falls under it;
  - a broader or more general category of which the subject is a significant
    kind, unless that category is so broad the subject is only one of many
    unrelated members;
  - a grouped label that explicitly names the subject as one of its components.

Otherwise REJECT. Relatedness is not identity: reject a candidate that is merely
associated or co-occurring with the subject, that the subject produces or gives
rise to, that is a downstream consequence or product of it, or that is a
separate sibling or relative not itself falling under the subject. Default to
REJECT; when genuinely unsure, REJECT.

Answer with a one-sentence reason naming the relationship, then the boolean.
```

**User prompt template:**

```
Subject: {subject_anchor}
Kind: {subject_kind}
Candidate: {candidate_name}
```

**Output schema** (`subject_trust` field on the PairJudgment):

```
{
  "reasoning": "<str>",
  "belongs": <bool>,
  "subject_name": "<str>",
  "subject_kind": "<str>"
}
```

#### S14.4 Shared output schemas

##### S14.4.1 `EvidenceQuality`

Used by the pair evidence judge, cross-document judge, and region assessment agents. The `overall` score is probability-anchored: 1 (<5%) through 5 (~50%) to 9 (>95%).

```
{
  "directness":  "explicit" | "implied" | "tangential",
  "source_type": "primary" | "review" | "other",
  "specificity": "mechanistic" | "associative" | "vague",
  "language":    "definitive" | "hedged" | "speculative",
  "overall":     <1-9>
}
```

##### S14.4.2 `PaperQualityAssessment`

Used by the document analysis agent. Seven dimensions, each scored 0–4 with required justification (see Section S5 for the full rubric). Overall score: 0–28; tiers: 0–7 exclude, 8–14 caution, 15–21 acceptable, 22–28 high trust.

```
{
  "method_clarity":          {"score": <0-4>, "justification": "<str>"},
  "data_provenance":         {"score": <0-4>, "justification": "<str>"},
  "statistical_rigour":      {"score": <0-4>, "justification": "<str>"},
  "internal_consistency":    {"score": <0-4>, "justification": "<str>"},
  "plausibility":            {"score": <0-4>, "justification": "<str>"},
  "reproducibility_signals": {"score": <0-4>, "justification": "<str>"},
  "integrity_indicators":    {"score": <0-4>, "justification": "<str>"}
}
```

Marker 2.0: An Updated Database of Manually Curated Cell Markers in Human/Mouse and Web Tools Based on scRNA-seq Data." *Nucleic Acids Research* 51 (D1): D870–76. https://doi.org/10.1093/nar/gkac947.

Huang, Yuxuan, Yihang Chen, Haozheng Zhang, Kang Li, Meng Fang, Linyi Yang, Xiaoguang Li, et al. 2025. "Deep Research Agents: A Systematic Examination And Roadmap." June 22, 2025. https://doi.org/10.48550/arXiv.2506.18096.

Islamaj, Rezarta, Po-Ting Lai, Chih-Hsuan Wei, Ling Luo, Tiago Almeida, Richard A. A Jonker, Sofia I. R Conceição, et al. 2024. "The Overview of the BioRED (Biomedical Relation Extraction Dataset) Track at BioCreative VIII." *Database* 2024 (February): baae069. https://doi.org/10.1093/database/baae069.

Lai, Po-Ting, Chih-Hsuan Wei, Ling Luo, Qingyu Chen, and Zhiyong Lu. 2023. "BioREx: Improving Biomedical Relation Extraction by Leveraging Heterogeneous Datasets." *Arxiv*, June, arXiv:2306.11189v1. https://www.ncbi.nlm.nih.gov/pmc/articles/PMC10370213/.

Li, Bo, Gexiang Fang, Yang Yang, Quansen Wang, Wei Ye, Wen Zhao, and Shikun Zhang. 2023. "Evaluating ChatGPT's Information Extraction Capabilities: An Assessment of Performance, Explainability, Calibration, and Faithfulness." April 23, 2023. https://doi.org/10.48550/arXiv.2304.11633.

Liu, Peiwen, Sakura Eri B Maezono, Weitao Lin, Yen Yeow, Ya-Yu Liu, Yasmin Hisham, Rui Hou, Jordan A Ramilowski, and Alistair R R Forrest. 2026. "connectomeDB2025: A Rigorously Curated, Multi-Species Resource of Experimentally Supported Ligand–Receptor Interactions." *Nucleic Acids Research* 54 (D1): D546–54. https://doi.org/10.1093/nar/gkaf1108.

Mayers, Michael, Tong Shu Li, Núria Queralt-Rosinach, and Andrew I. Su. 2019. "Time-Resolved Evaluation of Compound Repositioning Predictions on a Text-Mined Knowledge Network." *Bmc Bioinformatics* 20 (1): 653. https://doi.org/10.1186/s12859-019-3297-0.

Moreau, Erwan. 2023. "Literature-Based Discovery: Addressing the Issue of the Subpar Evaluation Methodology." *Bioinformatics* 39 (2): btad090. https://doi.org/10.1093/bioinformatics/btad090.

Nazi, Zabir Al, Vagelis Hristidis, Aaron Lawson McLean, Jannat Ara Meem, and Md Taukir Azam Chowdhury. 2025. "Ontology-Guided Query Expansion for Biomedical Document Retrieval Using Large Language Models." arXiv.org. August 15, 2025. https://arxiv.org/abs/2508.11784v1.

Oliveira, Rodrigo de, Matthew Garber, James M Gwinnutt, Emaan Rashidi, Jwu-Hsuan (Shantina) Hwang, William Gilmour, Jay Nanavati, Khaldoun Zine El Abidine, and Christina DeFilippo Mack. 2025. "A Study of Calibration as a Measurement of Trustworthiness of Large Language Models in Biomedical Natural Language Processing." *Jamia Open* 8 (4): ooaf058. https://doi.org/10.1093/jamiaopen/ooaf058.

Pandit, Shrey, Jiawei Xu, Junyuan Hong, Zhangyang Wang, Tianlong Chen, Kaidi Xu, and Ying Ding. 2025. "MedHallu: A Comprehensive Benchmark for Detecting Medical Hallucinations in Large

Language Models." February 20, 2025. https://doi.org/10.48550/arXiv.2502.14302.

Ramilowski, Jordan A., Tatyana Goldberg, Jayson Harshbarger, Edda Kloppmann, Marina Lizio, Venkata P. Satagopam, Masayoshi Itoh, et al. 2015. "A Draft Network of Ligand–Receptor-Mediated Multicellular Signalling in Human." *Nature Communications* 6 (1): 7866. https://doi.org/10.1038/ncomms8866.

Seo, Jean, Jongwon Lim, Dongjun Jang, and Hyopil Shin. 2024. "DAHL: Domain-specific Automated Hallucination Evaluation of Long-Form Text through a Benchmark Dataset in Biomedicine." November 14, 2024. https://doi.org/10.48550/arXiv.2411.09255.

Smith, Nathaniel, Xinyu Yuan, Chesney Melissinos, and Gaurav Moghe. 2025. "FuncFetch: An LLM-assisted Workflow Enables Mining Thousands of Enzyme–Substrate Interactions from Published Manuscripts." *Bioinformatics* 41 (1): btae756. https://doi.org/10.1093/bioinformatics/btae756.

Swanson, Don R. 1986. "Undiscovered Public Knowledge." *The Library Quarterly: Information, Community, Policy* 56 (2): 103–18. https://doi.org/10.1086/601720.

Thilakaratne, Menasha, Katrina Falkner, and Thushari Atapattu. 2019. "A Systematic Review on Literature-Based Discovery Workflow." *Peerj Computer Science* 5 (November): e235. https://doi.org/10.7717/peerj-cs.235.
